# Electrophysiological Indicators of Sleep-associated Memory Consolidation in 5- to 6-Year-Old Children

**DOI:** 10.1101/2020.09.04.283606

**Authors:** Ann-Kathrin Joechner, Sarah Wehmeier, Markus Werkle-Bergner

**Affiliations:** Center for Lifespan Psychology, Max Planck Institute for Human Development, Berlin, Germany

**Author notes:** Correspondence concerning this article should be addressed to: Ann-Kathrin Joechner or Markus Werkle-Bergner. Author Contributions:* A-KJ, SW, and MW-B designed the study; A-KJ and SW performed the experiments; A-KJ and MW-B analysed the data; A-KJ and MW-B wrote the manuscript. All authors revised the manuscript. Data Availability:* The custom code and data necessary to reproduce the results in this manuscript will be made available after acceptance of the manuscript on OSF https://osf.io/7mf9v/. During review, data and code are available from the corresponding authors upon request. Author Notes:* This research was conducted within the project “Lifespan Rhythms of Memory and Cognition (RHYME)” at the Max Planck Institute for Human Development. A-KJ is a fellow of the International Max Planck Research School on the Life Course (LIFE; http://www.imprs-life.mpg.de). MW-B received support from the German Research Foundation (DFG, WE 4269/5-1) and the Jacobs Foundation (Early Career Research Fellowship 2017–2019). We would like to thank B. E. Muehlroth for constant advice and for providing us with her analysis pipeline and Julia Delius for editorial assistance. We are grateful to the members of the RHYME and LIME projects for helpful feedback on the analyses. We further thank all the children and their families who participated in this study. Ethics Approval Statement:* The study was designed in agreement with the Declaration of Helsinki and was approved by the local ethics committee of the Max Planck Institute for Human Development. Permission to Reproduce Material from Other Sources:* Not applicable.

## Abstract

In adults, the synchronised interplay of sleep spindles (SP) and slow oscillations (SO) supports memory consolidation. Given tremendous developmental changes in SP and SO morphology, it remains elusive whether across childhood the same mechanisms as identified in adults are functional. Based on topography and frequency, we characterise slow and fast SPs and their temporal coupling to SOs in 24 pre-school children. Further, we ask whether slow and fast SPs and their modulation during SOs are associated with behavioural indicators of declarative memory consolidation as suggested by the literature on adults. Employing an individually tailored approach, we reliably identify an inherent, development-specific fast centro-parietal SP type, nested in the adult-like slow SP frequency range, along with a dominant slow frontal SP type. Further, we provide evidence that the modulation of fast centro-parietal SPs during SOs is already present in pre-school children. However, the temporal coordination between fast centro-parietal SPs and SOs is weaker and less precise than expected from research on adults. While we do not find evidence for a critical contribution of SP–SO coupling for memory consolidation, crucially, slow frontal and fast centro-parietal SPs are each differentially related to sleep-associated consolidation of items of varying quality. While a higher number of slow frontal SPs is associated with stronger maintenance of medium-quality memories, a higher number of fast centro-parietal SPs is linked to a greater gain of low-quality items. Our results demonstrate two functionally relevant inherent SP types in pre-school children although SP–SO coupling is not yet fully mature.

## 1 Introduction

It is widely agreed that rhythmic neuronal activity during sleep supports declarative memory consolidation (Diekelmann & Born, 2010; Watson & Buzsáki, 2015). Compelling evidence suggests that the underlying key mechanism is the reactivation of initially labile, learning-related, neuronal activity in the hippocampus and its integration into cortical networks (Peyrache et al., 2009; Wilson & McNaughton, 1994). This system consolidation results in more durable and integrated mnemonic representations (Diekelmann & Born, 2010). However, given profound developmental changes in rhythmic neuronal activity, it is still unclear whether the neuronal mechanisms facilitating sleep-associated system consolidation identified in young adults apply similarly to children of all ages.

The canonical view suggests that system consolidation mainly takes place during non-rapid eye movement sleep (NREM) through precise temporal co-occurrence of hippocampal activity with fast sleep spindles (SPs, ≍12–16 Hz), initiated by the UP state of the slow oscillation (SO, < 1 Hz, Buzsáki, 1998; Clemens et al., 2007; Diekelmann & Born, 2010; Helfrich et al., 2018; Klinzing et al., 2016; Latchoumane et al., 2017; Mölle et al., 2009, 2011). SOs are marked by high-amplitude UP and DOWN states, reflecting changes in the membrane potential of large populations of cortical neurons alternating between joint depolarisation and hyperpolarisation, respectively (Steriade, Contreras et al., 1993; Steriade, Nunez, & Amizica, 1993). Via cortico-thalamic pathways, SO UP state depolarisation creates conditions in the thalamus that initiate SPs (Contreras et al., 1997; Steriade, 2006). SPs arise through reciprocal interactions between reticular thalamic and thalamo-cortical neurons and the latter transmit them to the cortex where they are thought to induce increased plasticity (Bonjean et al., 2011; Lüthi, 2014; Muller et al., 2016; Niethard et al., 2018; Rosanova & Ulrich, 2005; Timofeev et al., 2002). Fast SPs, in turn, have been repeatedly shown to orchestrate hippocampal activity and to facilitate hippocampal-cortical connectivity in rodents and humans (Andrade et al., 2011; Clemens et al., 2007; Siapas & Wilson, 1998; Sirota et al., 2003). Thus, fast SPs offer perfect conditions for the integration of hippocampal activity patterns into cortical networks (Muller et al., 2016; Niethard et al., 2018). Importantly, while fast SPs coupled with hippocampal activity can also occur independently of SOs, it is the triad of SOs, fast SPs, and hippocampal activity that seems to be beneficial for memory (Helfrich et al., 2018; Latchoumane et al., 2017; Muehlroth et al., 2019; Nir et al., 2011).

Besides a fast SP type, predominant in centro-parietal brain areas, there is also a slow, frontal SP type (≍9–12 Hz, Andrillon et al., 2011; Mölle et al., 2011) identified in surface electroencephalography (EEG) in adults. Slow SPs differ from the fast type in several aspects including, e.g., frequency, topography, circadian regulation, preferred phase of occurrence during SOs, their expression across the lifespan, and role for memory (De Gennaro & Ferrara, 2003; Fernandez & Lüthi, 2020). That is, contrary to fast SPs, slow SPs are less numerous during the UP state, but occur more often during the transition from the UP to the DOWN state. Furthermore, their role for system consolidation is less established. However, it has been hypothesised that rather than hippocampal-neocortical integration, slow SPs may be preferentially involved in cortico-cortical storage mechanisms (Astori et al., 2013; Ayoub et al., 2013; Doran, 2003; Rasch & Born, 2013; Timofeev & Chauvette, 2013).

During development, rhythmic neuronal activity patterns change drastically in their temporal expression, peak frequency, and topographical distribution (Clarke et al., 2001; Hahn et al., 2019; Purcell et al., 2017). The hallmark of the presence of a given rhythm is the existence of an identifiable peak in the power spectrum (Aru et al., 2015; Kosciessa et al., 2020). Considering the adult pattern as a point of reference (slow SPs: 9–12 Hz, frontal distribution; fast SPs: 12–16 Hz, centro-parietal distribution; SOs: < 1 Hz, frontal distribution), the following age-differences were reported in developmental samples. Slow rhythmic neuronal activity (0.5–4 Hz), comprising SOs, is most strongly pronounced before the onset of puberty, attenuating thereafter, that is, in SO power and steepness of the slope (Campbell & Feinberg, 2009; Kurth, Jenni et al., 2010). Moreover, in contrast to SOs in young adults that predominantly originate in anterior cortical regions, SO onset and spectral dominance was located in central and posterior areas respectively in pre-pubertal children (Kurth, Ringli et al., 2010; Timofeev et al., 2020).

Unlike SOs, overall, SPs are observed increasingly often between the age of 4 until early adulthood (Olbrich et al., 2017; Purcell et al., 2017; Scholle et al., 2007). Systematic research on the differentiation between slow and fast SPs in children has been scarce so far. Whereas adult-like slow SPs already evolve early in childhood, adult-like fast SP development seems to be prolonged (D’Atri et al., 2018; Hahn et al., 2019; Purcell et al., 2017). Around the age of 4–5 years, adult-like fast SPs are comparatively rare and only fully emerge around puberty several years later (D’Atri et al., 2018; Hahn et al., 2019; Purcell et al., 2017). In addition, it is typically challenging to provide evidence for separate SP types in children. The gold standard for identification of more than one rhythm in the SP range in children would require evidence of two separate spectral peaks (Aru et al., 2015; Kosciessa et al., 2020). Independent of recording site, SP peaks usually fall within the adult-like slow SP frequency range, suggesting a dominance of adult-like slow SPs during childhood (Hoedlmoser et al., 2014; Shinomiya et al., 1999). Still, early descriptive research points towards a possible dichotomy of SP types based on frequency and topography already around the age of 2 years (Jankel & Niedermeyer, 1985; Shinomiya et al., 1999). Across maturation, the peak frequency increases with an adult-like fast centro-parietal peak evolving around puberty (Campbell & Feinberg, 2009; Shinomiya et al., 1999; Tarokh & Carskadon, 2010).

In line with findings of less pronounced SPs across childhood, a recent study also reported lower coupling between SPs and SOs during childhood that increased across adolescence (Hahn et al., 2020). Together, the existing literature suggests that the assumed core mechanisms of sleep-associated memory consolidation, i.e., adult-like fast SPs and the temporal synchronisation of SPs by SOs, might not yet be fully functional in children.

However, there is no reason to assume a priori that functionally equivalent rhythmic neuronal events are expressed in exactly the same way across the lifespan (Clarke et al., 2001; Olbrich et al., 2017; Shinomiya et al., 1999). Nevertheless, analyses of sleep electrophysiology in children mostly rely on the application of fixed adult-derived (nevertheless often inconsistent) criteria without ensuring the presence of a rhythm (i.e., a spectral peak) within the search space (but see Friedrich et al., 2019; Olbrich et al., 2017). As a result, it often remains elusive whether the rhythmic neuronal phenomenon of interest in a given child during a given developmental period is reliably captured or whether functionally different rhythmic neuronal events might be mixed (Cox et al., 2017; Ujma et al., 2015). Considering evidence from adults for distinct functions, this applies specifically to slow and fast SPs during childhood. A distinction into slow and fast SPs based on adult criteria may simply miss relevant developmental shifts. Therefore, it is unclear whether the scarce findings on fast SPs in children indeed reflect a missing fast SP rhythm or a bias of the analysis approach. Given well-known developmental frequency acceleration (Campbell & Feinberg, 2016; Marshall et al., 2002), it is conceivable that a functionally relevant, development-specific fast SP type might already be present in children, though expressed at slower frequencies than in adults. Hidden in the adult slow-frequency range, this may hamper the fast type’s identification (Olbrich et al., 2017). Thus, imprecisely capturing the within-person, age-specific neuronal rhythm of interest may pose specific challenges when aiming to uncover its mechanistic role in memory consolidation across development (Muehlroth & Werkle-Bergner, 2020).

Indeed, while numerous studies confirm the importance of sleep for declarative memories across childhood, evidence on the electrophysiological correlates of system consolidation mechanisms during sleep remains scarce and inconsistent (Ashworth et al., 2014; Backhaus et al., 2008; Friedrich et al., 2019, 2020; Hahn et al., 2019; Hoedlmoser et al., 2014; Kurdziel et al., 2013; Wilhelm et al., 2008, 2013, 2020). Comparable to findings in adults, both SPs and slow neuronal activity have been related to sleep-associated memory consolidation in children; but notably with inconsistencies across memory tasks and age ranges (Friedrich et al., 2015, 2019; Hahn et al., 2019; Hoedlmoser et al., 2014; Kurdziel et al., 2013; Maski et al., 2015; Prehn-Kristensen et al., 2011, 2014; Wang et al., 2017). Part of these inconsistencies may result from the simple extrapolation of adult-derived criteria for the detection of sleep-associated neuronal rhythms. While most studies did not differentiate between slow and fast SPs, recent longitudinal findings suggest that the development of adult-like fast SPs and enhanced temporal synchrony between SPs and SOs supports effects of sleep on memory from pre-pubertal childhood to adolescence (Hahn et al., 2019; Hahn et al., 2020). Precise and development-sensitive detection of neuronal rhythms may therefore benefit the reliable identification of electrophysiological markers of sleep-associated memory consolidation in children.

Another source of inconsistencies across studies may result from ignoring the encoding strength of individual memories (i.e., memory quality; Craik & Lockhart, 1972; Tulving, 1967) prior to sleep (Muehlroth et al., 2020; Wilhelm et al., 2012, 2020). Memory quality has been suggested to impact the underlying processes of memory consolidation and their subsequent outcomes — that is, either memories already accessible prior to sleep are maintained or items previously not consciously available are gained (Dumay, 2016, 2018; Fenn & Hambrick, 2013). Several studies have indicated that sleep-associated system consolidation mechanisms preferentially act on the maintenance of memories from weak to intermediate quality (Denis et al., 2020; Drosopoulos et al., 2007; Fenn & Hambrick, 2013; Muehlroth et al., 2020; Schapiro et al., 2018; Schreiner & Rasch, 2018; Wilhelm et al., 2012; but see Schoch et al., 2017; Tucker & Fishbein, 2008). Relying on memory performance averaged across items may thus introduce further unwanted noise (Tulving, 1967) when trying to disentangle the functions of sleep oscillations for memory consolidation across development. Hence, examining how sleep supports memory consolidation in childhood necessitates appropriate assessment of the electrophysiological and memory processes involved.

The present study targeted two main questions: Firstly, we asked whether the specific electrophysiological indicators of sleep-associated memory consolidation can be detected reliably in pre-school children. Therefore, we set out to characterise slow and fast SPs and their temporal interaction with SOs using individualised rhythm detection in pre-schoolers. Secondly, we asked whether SPs and their temporal modulation by SOs in 5- to 6-year-olds would show comparable associations with the behavioural indicators of memory consolidation as suggested by findings in adults. To control for inter-individual differences in memory quality, we adapted a paradigm developed to control for memory quality of individual items in adults (Fandakova et al., 2018; Muehlroth et al., 2019) for use in pre-school children.

## 2 Methods

### 2.1 Participants

Thirty-six pre-school children (19 female, *M_age_*= 69.53 mo, *SD_age_* = 6.50 mo) were initially enlisted to participate in our exploratory study on the role of sleep oscillations in memory consolidation in pre-school children. Participants were recruited from daycare centres in Berlin, Germany and from the database of the Max Planck Institute for Human Development (MPIB). Five participants did not complete the study protocol. Data collection from four children was incomplete due to technical failures during one of the two polysomnographic (PSG) recordings. Additionally, three participants were excluded from further analyses because they failed to complete the behavioural task. Therefore, the final sample consisted of 24 children (14 females; *M*_age_ = 70.71, *SD_age_* = 7.28 mo). The participants were randomly assigned to one of two learning conditions: (1) the children studied 50 scene-object associations (*N* = 14, *M_age_* = 68.57, *SD_age_* = 7.51 mo) and (2) the children studied 100 scene-object associations (*N* = 10, *M_age_* = 73.70, *SD*_age_ = 6.08 mo). The two groups did not differ significantly in their mean age (*Z* = −1.76, *p* = 0.078, *CI*_2.5, 97.5_ [−2.44; 0.00]). All participants were native German speakers without current or chronic illness, use of medication, personal or family history of mental or sleep disorders, obesity (body mass index > 28 kg/m²), respiratory problems (e.g., asthma), and without evidence of a learning disability. All participants completed a short screening prior to study participation. Subjective sleep quality was assessed by the Children’s Sleep Habit Questionnaire (CSHQ, Schlarb et al., 2010) and the Children’s Sleep Comic (SCC, Schwerdtle et al., 2012). The Strengths and Difficulties Questionnaire (SDC, Goodman, 1997) was used to screen for behavioural and emotional difficulties. In addition, parents filled in the Children’s Chronotype Questionnaire (Werner et al., 2009), a short demographic questionnaire, and a sleep log starting three days before the first PSG night. Children received a gift for their participation and their families received monetary compensation. The study was designed in agreement with the Declaration of Helsinki and was approved by the local ethics committee of the MPIB.

### 2.2 General Procedure

The experimental protocol for each participant encompassed seven days and included two nights of electrophysiological sleep recordings (Figure 1). Sleep was recorded in the participants’ familiar environment using ambulatory PSG (SOMNOscreen plus, SOMNOmedics GmbH, Randersacker, Germany). PSG recordings started and ended corresponding to each participant’s individual bedtime habits. The first night served as an adaption and baseline night (Figure 1, baseline night). The second was flanked by an associative scene-object memory task with cued recall before and after sleep (Figure 1, learning night; for details, see below). We contrasted indicators of sleep quality (see “2.5.2 EEG Pre-processing”) between the two nights and found no differences between baseline and learning night (Supplementary Table S1). All behavioural assessments took place in a standardised laboratory environment at the MPIB. Three days before the first night, children’s sleep was stabilised according to their habitual bed and wake times and monitored by sleep logs filled in by the parents together with their children.

**Figure 1.**
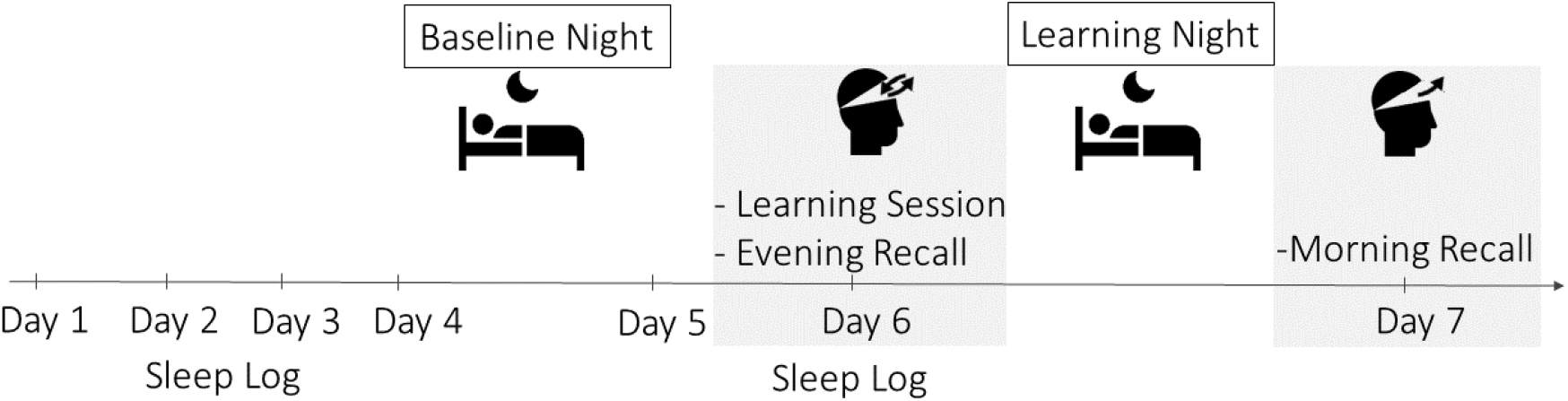
Experimental Procedure. Sleep was monitored for two nights (baseline and learning night) using ambulatory PSG. Participants started filling in a daily sleep log three days before the baseline night and continued throughout the whole procedure. The memory task took place before and after the learning night (grey boxes).

### 2.3 Memory Task

The memory task used in the present study was a child-adapted version of an associative scene-word memory paradigm designed to trace the quality of associative memories within individuals using repeated cued recall sessions (Fandakova et al., 2018; Muehlroth et al., 2019). In order to adapt the task to our population of 5- to 6-year-old children who were just beginning to learn to read, we replaced the written nouns with photographs of everyday objects. To make the task more appealing, children were told that they would play a game similar to the “Memory” game.

Preceding the main memory paradigm, participants were encouraged to try to remember a scene-object pair by integrating the scene and the object into one joint vivid mental image. Participants trained use of this child-appropriate imagery strategy (e.g., Danner & Taylor, 1973) in 10 trials that were not part of the main task. For the first example, the experimenter would explain step by step what it means to create one joint mental image in child-friendly language. During this procedure, the child was shown a joint image where the object had actually been placed into the scene. In addition, children were given tips on how to make such a joint image vivid. For the next three trials, the child was asked to try out creating a joint mental image that they found really funny or unusual and then to verbalise it. Independent of the answer, they were presented with an example of what one could imagine for that specific scene-object pair. They continued training with another six trials in which they were not presented with an example afterwards. The learning strategy training took 15 minutes. It was implemented to minimise age-related and inter-individual differences in strategy acquisition and usage during the subsequent main task (Schneider & Sodian, 1997; Shing et al., 2008).

During the *encoding phase* (encoding, Figure 2), scene-object pairs were presented on a screen for 4000 ms. Each pair was followed by the presentation of a 3-point Likert scale for 2000 ms where participants were asked to indicate how well they were able to apply a previously trained imagery strategy (see above). A fixation cross shown for 1000 ms separated individual trials. Immediately after the encoding session, a *cued recall plus feedback session* followed (recall & feedback, Figure 2). The scenes served as cues and participants were asked to verbally recall the object corresponding to the scene within a 8000 ms interval. For this purpose, the object to be recalled was covered by the back of a playing card on the screen. The correctness of answers was coded by the experimenter and also audio-recorded. Subsequently, the correct pairing was again presented for 2000 ms irrespective of the previous answer. This feedback was intended to provide an additional learning opportunity. The availability of specific scene-object associations before and after sleep was tested by a cued recall test prior to bedtime (*evening recall*, Figure 2) and during a cued recall test in the morning (*morning recall*, Figure 2). The evening recall took place in the evening following a 10 min break after the cued recall plus feedback session. The morning recall was performed in the morning, two hours after the participant woke up.

**Figure 2.**
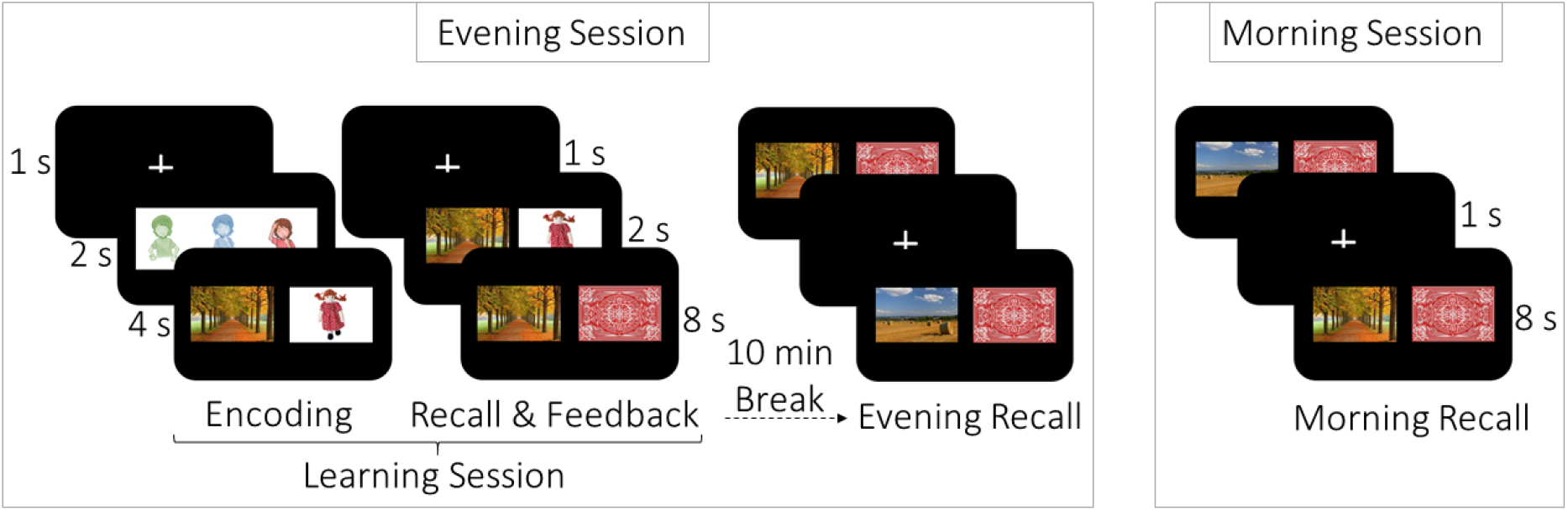
Associative Memory Task. During the evening session, participants studied scene-object pairs in two runs (learning session; encoding, recall & feedback) and were tested for their memory of the learned pairs during the evening recall. In the morning, participants were again probed for their memory of the learned scene-object pairs with a cued recall task (morning session; morning recall).

To further adapt task procedures for young children, we reduced the original number of stimuli to adjust task difficulty and attention requirements. Due to a lack of comparable studies in this age group, but based on similar studies in older children (Hoedlmoser et al., 2014; Urbain et al., 2016), we created two lists of different trial lengths: one with 50 and another with 100 non-associated pairs of scenes and objects. This resulted in two groups learning a different number of scene-object associations (henceforth labelled Group_50_ and Group_100_). As we could not determine the appropriate number of scene-object pairs for 5- to 6-year-old children a priori, the trial length manipulation was initially intended to explore the task-difficulty space in this age range. However, as it turned out in the analyses, trial-length groups did not differ with regard to their memory performance (Supplementary Table S2, Supplementary Figure S1). Hence, for most analyses, we collapsed across both groups. Nevertheless, we indicate group membership in result plots. Memory performance showed neither ceiling nor floor effects (Supplementary Figure S1).

The whole learning session lasted no more than 30 min (Group_100_, including breaks). Pairs of scenes and objects were presented on a black background on a 15.6’’ screen. Scenes were always displayed in the left hemifield and objects in the right hemifield. The order of presentation was randomised across learning and cued recall sessions but not across participants. In addition, the first 50 trials were equal for Group_50_ and Group_100_. The task was implemented using PsychToolbox (Kleiner et al., 2007) for Matlab (MathWorks, Natick, MA).

### 2.4 Behavioural Analyses

General performance during recall sessions was calculated as the ratio of correctly recalled objects to the total number of trials (i.e., 50 or 100) multiplied by 100. The effect of sleep on memory consolidation was determined as the probability to successfully recall an item during the morning recall. Given the two recall sessions in the evening (recall & feedback, evening recall), we were able to analyse the effect of sleep on memory contingent on each item’s recall history during the evening session (Figure 2, Dumay, 2016; Muehlroth et al., 2020). Based on an item’s individual retrieval success during the evening, we distinguished three categories of memory quality (Tulving, 1967) and assessed the effect of sleep on memory consolidation within these three categories separately. Firstly, items not recalled during either of the two evening recalls (recall & feedback and evening recall) were categorised as items of *low memory quality*. Secondly, items correctly remembered during both evening recalls were categorised as items of *high memory quality*. Finally, items recalled correctly only once during the evening session were considered as being of *medium memory quality*. In other words, these were items remembered during the evening recall but not during the recall & feedback session as well as items remembered during the recall & feedback session but not during the evening recall. Even though theoretically less likely, we decided to include the latter scenario in our analyses because successful recall during the recall & feedback session indicates that an accessible memory trace was initially established, even if it was not remembered in the evening recall. In itself, the lack of successful retrieval during the evening recall is not indicative of a complete deterioration of this memory trace (Tulving & Pearlstone, 1966; Tulving & Psotka, 1971). Such items are likely still available in the memory system for memory consolidation to act upon, they may just be temporarily inaccessible (Habib & Nyberg, 2008). In fact, the ability to recall an item during the recall & feedback session but not during the evening recall was evident in 13 out 24 participants and applied to only a small number of items except for one subject from Group_100_ where it applied to 39 items.

Our main behavioural analyses are centred on items successfully retrieved during the morning recall for the different memory quality levels. In this context, low-quality items (i.e., those that were not recalled at all during the evening session) that were successfully retrieved during morning recall can be regarded as gained items. By contrast, morning recall of medium- and high-quality items (i.e., those that were recalled at least once during the evening session) reflects memory maintenance (Dumay, 2016; Fenn & Hambrick, 2013; Muehlroth et al., 2020).

### 2.5 Sleep Polysomnography Acquisition and Analyses

#### 2.5.1 Data Acquisition

Sleep was recorded using an ambulatory PSG system (SOMNOscreen plus, SOMNOmedics GmbH, Randersacker, Germany). For the EEG recordings, a total of 15 gold electrodes were positioned according to the international 10-20 System (Jasper, 1958) including left and right horizontal electrooculogram (HEOG), two submental electrodes referenced against one chin electrode for electromyogram (EMG), and seven active scalp electrodes (F3, F4, C3, Cz, C4, Pz, Oz). The ground electrode was placed at AFz. Two electrodes were placed on the left and right mastoids (A1, A2) for later re-referencing. The EEG data were recorded between 0.2 and 75 Hz at a sampling rate of 128 Hz against the common reference Cz. In addition, cardiac activity was recorded using two electrocardiogram (ECG) derivations. Impedances were kept below 6 kΩ, prior to start of the recordings.

#### 2.5.2 EEG Pre-processing

Initially, PSG data were offline filtered and re-referenced against the averaged mastoids (A1, A2) for visual sleep stage identification using BrainVisionAnalyzer 2.1 (Brain Products GmbH, Gilching, Germany). Two scorers then visually classified sleep stages in epochs of 30 s according to the rules of the American Academy of Sleep Medicine (Berry et al., 2015) using the program SchlafAus 1.4 (Copyright © by S. Gais & M. Werner, 2006). Based on the visual scoring, the following indicators of sleep quality (Ohayon et al., 2017) were calculated: (1) total sleep time (TST, the time spent in N1, N2, N3, and R), (2) percentage N1, N2, N3, and R (the time spent in a respective sleep stage relative to TST), and (3) wake after sleep onset (WASO, the time awake between sleep onset and final awakening). Afterwards, using Matlab R2016b (Mathworks Inc., Sherbom, MA) and the Fieldtrip toolbox (Oostenveld et al., 2011), the EEG data were semi-automatically cleaned for the detection of rhythmic neuronal events during sleep. In a first step, bad EEG channels were rejected based on visual inspection. Then, an automatic artefact detection algorithm was implemented for the remaining channels on 1 s epochs to exclude segments with strong deviations from the overall amplitude distribution. Therefore, mean amplitude differences were Z-standardised within each segment and channel. Segments were marked as bad if either visually identified as body movements or if they exceeded an amplitude difference of 500 µV. Furthermore, segments with a *Z*-score exceeding 5 in any channel were excluded (see Muehlroth et al., 2019 for similar procedures).

#### 2.5.3 Detection of Rhythmic Neuronal Activity

##### 2.5.3.1 Sleep Spindle Detection

SPs were detected during NREM sleep (N2 and N3) using an established algorithm (Klinzing et al., 2016; Mölle et al., 2011; Muehlroth et al., 2019; see Supplementary Figure S2 for raw EEG traces with SPs) with individually adjusted frequency bands and amplitude thresholds (Muehlroth & Werkle-Bergner, 2020). SPs vary considerably across individuals (Cox et al., 2017; Ujma et al., 2015; Werth et al., 1997) and development (Hahn et al., 2019; Purcell et al., 2017). Accordingly, previous studies have shown that individualised SP detection approaches yield the most sensitive detection results (Adamczyk, 2015; Ujma et al., 2015). Since a slow SP type has been shown to be more prevalent in frontal areas and a fast SP type to be predominant in central and parietal areas (Anderer et al., 2001; De Gennaro & Ferrara, 2003), we firstly identified the individual SP peak frequency in the 9–16 Hz range in averaged frontal and centro-parietal electrodes (Ujma et al., 2015). Power spectra were calculated for NREM sleep during the baseline and the learning night, respectively, by applying a Fast-Fourier Transform (FFT) on every 5 s artefact-free epoch using a Hanning taper. On the assumption that the EEG background spectrum is characterised as A*f^-a^ (Buzsáki & Mizuseki, 2014), the resulting power spectra were then fitted linearly in the log (frequency)-log (power) space using robust regression to model the background spectrum. The estimated background spectrum was then subtracted from the original power spectrum (Figure 3A, B). Using this approach, the resulting peaks in the power spectrum represent rhythmic, oscillatory activity (Kosciessa et al., 2020). Finally, the frontal and centro-parietal peak frequency was identified in the corrected power spectra with an automated algorithm combining a first derivative approach (Grandy et al., 2013) with a classical search for maxima (Supplementary Figure S3 for all individual power spectra). Individual frequency bands for SP detection in frontal, central, and parietal electrodes were defined as frontal or centro-parietal peak frequency ± 1.5 Hz, respectively (Mölle et al., 2011). EEG data were then band-pass filtered using a Butterworth two-pass filter of 6^th^ order for the respective frequency bands and the root mean square (RMS) was calculated at every sample point using a sliding window of 0.2 s. The resulting RMS signal was smoothed with a moving average of 0.2 s. SPs were detected whenever the amplitude of the RMS signal exceeded the mean of the filtered signal by 1.5 SD for 0.5–3 s. Successive SPs with boundaries within an interval of 0.25 s were merged if the resulting event did not exceed 3 s. Within such a merging run, one SP could only be merged with one other SP. The merging process was repeated iteratively until no further merging was possible (Mölle et al., 2011; Muehlroth et al., 2019). Only SP events detected in artefact-free segments were considered. Given the results of our time-frequency analyses of the temporal association between SPs and SOs, we additionally extracted SPs in frontal, central, and parietal electrodes higher than the individually identified upper limit for the event coupling analyses (henceforth called “high” SPs; peak frequency + 1.5 Hz < “high” SPs < 16 Hz; see Figure 3B, Supplementary Table S3 for descriptive measures of individually identified and “high” SPs).

**Figure 3.**
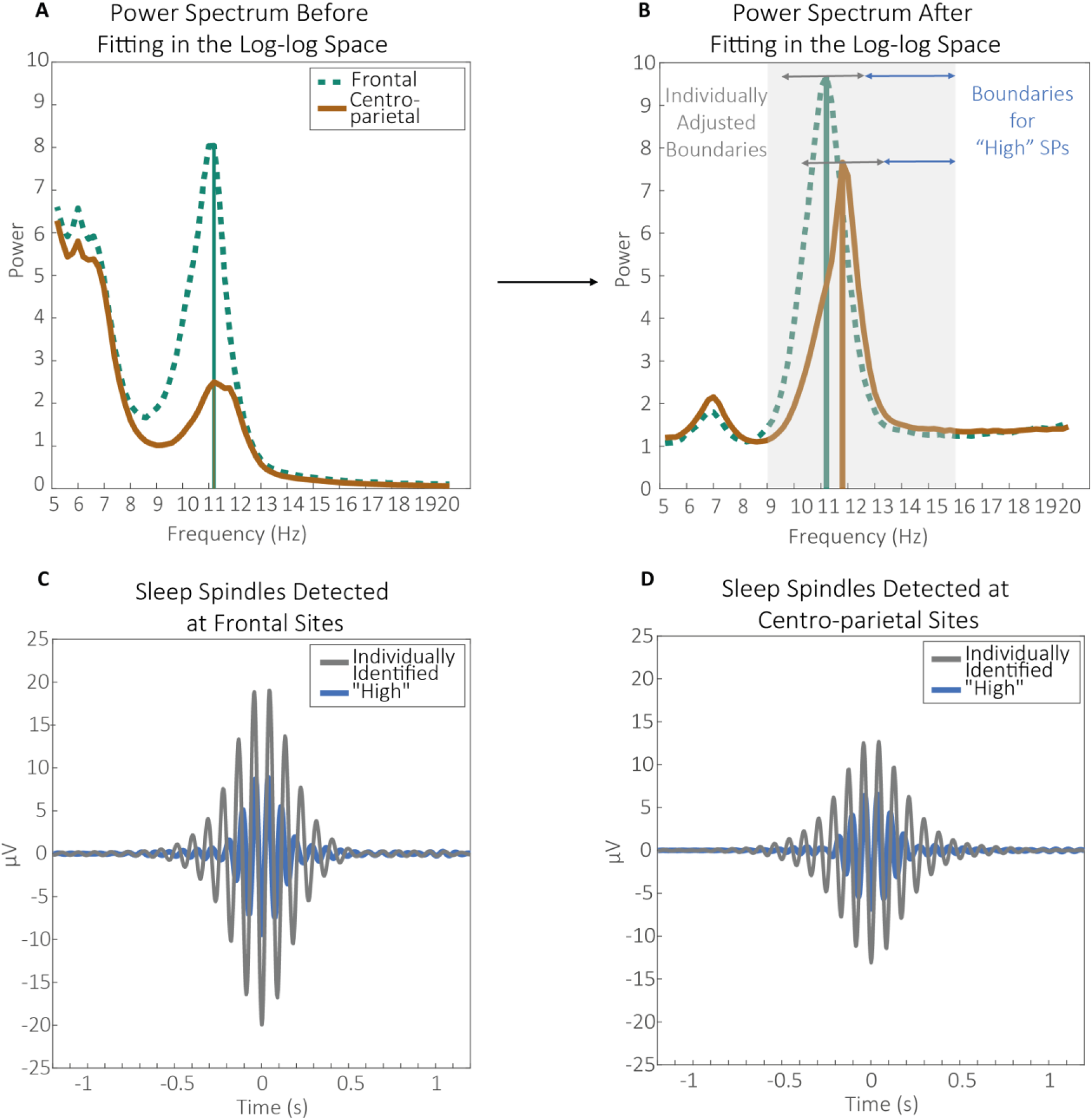
Schematic of the approach to define SP frequency boundaries based on the individual peak frequency and “high” SPs in averaged frontal and centro-parietal electrodes. (A) The original power spectra were (B) corrected by their background spectra. Based on the first derivative and the maximum peak, SPs were individually identified within ±1.5 Hz around the respective peaks. We additionally extracted SPs specifically higher than our individually identified upper boundary ((B) “High” SPs) for coupling analyses between SPs and SOs. (C & D) Shapes of averaged individually identified and “high” SPs detected in (C) frontal and (D) centro-parietal sites.

##### 2.5.3.2 Slow Oscillation Detection

Given the evidence that slow rhythmic neuronal activity during sleep shows a posterior rather than an anterior prevalence in children (Kurth, Ringli et al., 2010), SOs were detected at all electrodes during NREM sleep (N2 and N3, see Supplementary Figure S2 for raw EEG traces depicting SOs). Detection was based on Mölle et al. (2011) and Muehlroth et al. (2019) using an individualised amplitude criterion. The EEG signal was first filtered at 0.2–4 Hz using a Butterworth two-pass filter of 6^th^ order. Then, zero-crossings were detected in the filtered signal and positive and negative half-waves were identified. A negative half-wave, combined with a succeeding positive half-wave with a frequency of 0.5–1 Hz was considered a potential SO. A potential SO was finally considered a proper SO if its peak-to-peak amplitude exceeded 1.25 times the average peak-to-peak amplitude of all potential SOs and when the amplitude of the negative peak only exceeded 1.25 times the average negative amplitude of all putative SOs. Finally, only SOs that did not overlap with artefact segments were extracted.

#### 2.5.4 Statistical Analyses

Statistical analyses were conducted using the open-source toolbox Fieldtrip (Oostenveld et al., 2011) for Matlab (R2016b, Mathworks Inc., Sherbom, MA) and R 3.6.1 (R Core Team, 2014). Due to violations of the assumption of normality in some variables (Shapiro-Wilk Test) and given our small sample size, most analyses were based on non-parametric approaches. For correlations, pairwise-comparisons, and regression analyses we provide the 95% simple bootstrap percentile confidence interval (*CI*) of the respective parameter estimate, based on 5000 case re-samples. For regression analyses, we turned to robust methods whenever there were indicators for outliers, high leverage observations, or influential observations (QQ-Plot, Cook’s Distance). For repeated-measure Analyses of Variance (ANOVA), degrees of freedom were corrected according to Greenhouse-Geisser (*Ɛ* < 0.75) or Huynh-Feldt (*Ɛ*> 0.75) in cases of violations of sphericity. The generalised eta squared (*η^2^_G_*) is provided as a measure of effect size. Planned comparisons and post-hoc analyses for ANOVAs were conducted using non-parametric Wilcoxon rank-sum tests for independent comparisons and Wilcoxon signed-rank tests for dependent comparisons. Missing data was handled by listwise exclusion. All post-hoc tests were corrected for multiple comparisons using the Bonferroni-Holm method (Eichstaedt et al., 2013).

#### 2.5.5 Time-frequency Analyses of the Temporal Association Between Sleep Spindles and Slow Oscillations

To describe the modulation of SPs by SOs, a first set of analyses explored power modulations during SOs (Muehlroth et al., 2019). Analyses were conducted during NREM sleep (N2 and N3) on artefact-free segments only. Trials containing SOs were selected by centring the data ± 3 s around the DOWN peak of SOs. To allow for the interpretation of any SO-related power de- or increase, we matched every SO trial with a randomly chosen SO-free 6 s segment from the same electrode and sleep stage. Subsequently, time-frequency representations of 5–20 Hz were derived for trials with and without SOs using a Morlet wavelet decomposition (12 cycles) in steps of 1 Hz. Time-frequency representations during SO trials were then compared to SO-free trials within every participant using independent-sample *t*-tests. Given the high incidence of SOs during N3, we often identified a lower number of SO-free than of SO trials. To account for this, 100 random combinations of SO and SO-free trials were drawn and contrasts of power during SO trials versus SO-free trials were calculated for all 100 combinations and then averaged. The ratio of N2 to N3 trials was maintained during this procedure. The resulting *t*-maps represent power increases and decreases at 5–20 Hz during SO trials compared to SO-free trials within each individual. Finally, we conducted a cluster-based permutation test (Maris & Oostenveld, 2007) with 5000 permutations within a time segment of − 1.2–1.2 s (centred to the DOWN peak of the SO) to compare *t*-maps against zero on a group level. This time segment was chosen to cover one complete SO cycle (0.5–1 Hz, 1–2 s). We used a two-sided test with the critical alpha-level *α* = 0.05 (which means that each tail was tested with *α* = 0.025).

#### 2.5.6 Analyses of the Temporal Relation Between Discrete Sleep Spindles and Slow Oscillations

The general co-occurrence of discrete SPs and SOs was determined by identifying the percentage of SP centres (SO DOWN peaks) during NREM (N2 & N3) occurring within an interval of ± 1.2 s around the DOWN peak of SOs (SP centre), relative to all SPs (SOs) detected during NREM sleep. To explore the temporal coordination of SPs with respect to the SO cycle, we created peri-event time histograms (PETH) by determining the percentage of SP centres occurring within bins of 100 ms during an interval of ± 1.2 s around the SO DOWN peak. Percentage values within bins reflect the frequency of SP centres occurring within one bin relative to the total number of SP centres during the complete SO ± 1.2 s time interval (multiplied by 100). To test whether the occurrence of SPs within each bin was specific to the SO cycle as opposed to spontaneous occurrence, the individual percentage frequency distributions of SP centre occurrence were tested against surrogate distributions using dependent *t*-tests. The surrogate distributions were obtained separately for each individual by randomly shuffling the temporal order of the PETH bins 1000 times and then averaging across the 1000 sampling distributions. A cluster-based permutation test with 5000 permutations was applied to control for multiple comparisons. We used a two-sided test with the critical alpha-level *α* = 0.05 (which means that each tail was tested with *α* = 0.025).

## 3 Results

### 3.1 Individually Adjusted Sleep Spindle Detection Reveals Two Distinguishably Fast Sleep Spindle Types in Frontal and Centro-parietal Regions

Based on scarce evidence on the expression of fast SPs in children, we explored the possibility that children aged 5–6 years already express two inherent types of SPs: a slow frontal and fast centro-parietal SP type. Having established two distinguishable peaks in frontal and centro-parietal recording sites (Supplementary Figure S3), we then tested for evidence of two SP types by applying separate repeated-measure ANOVAs with the within-person factors NIGHT (baseline vs. learning) and ELECTRODE (F3, F4, C3, Cz, C4, Pz) on individually identified SP frequency, density, and amplitude during NREM (N2 & N3) sleep. Overall, none of the SP measures differed between nights (*F_frequency_*(1,18) = 0.27, *p* = 0.610, *η^2^_G_* < 0.01; *F_density_*(1,18) = 0.001, *p* = 0.970, *η^2^_G_* < 0.01; *F_amplitude_*(1,18) = 0.65, *p* = 0.431, *η^2^_G_* < 0.01). However, as expected, individually identified SPs varied overall in their frequency (*F*(1.18,21.28) = 32.68, *p* < 0.001, *η^2^_G_* = 0.33), density (*F*(2.92,52.55) = 14.98, *p* < 0.001, *η^2^_G_* = 0.18), and amplitude (*F*(1.53,27.53) = 53.41, *p* < 0.001, *η^2^_G_* = 0.42; Figure 4A–C) across electrodes. Planned contrasts comparing SP measures in frontal electrodes with central and parietal recording sites revealed significantly lower frequency in F3 and F4 as compared to C3, C4, Cz, and Pz (Figure 4, Supplementary Table S4A, all *Z* < − 5.00, all *p* < 0.001). Inversely, SP density and amplitude were significantly higher at frontal than at central and parietal recording sites (Figure 4, Supplementary Table S4A, all *Z* < −2.00, all *p* < 0.040). These effects did not differ between the baseline and learning night (interaction effects: *F_frequency_*(2.28,40.94) = 0.81, *p* = 0.465, *η^2^_G_* < 0.01; *F_density_*(2.58,46.49) = 1.07, *p* = 0.363, *η^2^_G_*< 0.01; *F_amplitude_*(2.72,48.88) = 0.71, *p* = 0.536, *η^2^_G_*< 0.01). Given the consistent differences between frontal and centro-parietal SP characteristics, we collapsed frontal (F3, F4) and centro-parietal (C3, Cz, C4, Pz) recording sites, creating measures representing a slower frontal and faster centro-parietal SP type for all following analyses. These two SP types are termed “slow frontal” and “fast centro-parietal” in the following. Since there were no differences in SP characteristics between the two nights, we focus on the learning night for all following analyses involving SPs, unless stated otherwise.

**Figure 4.**
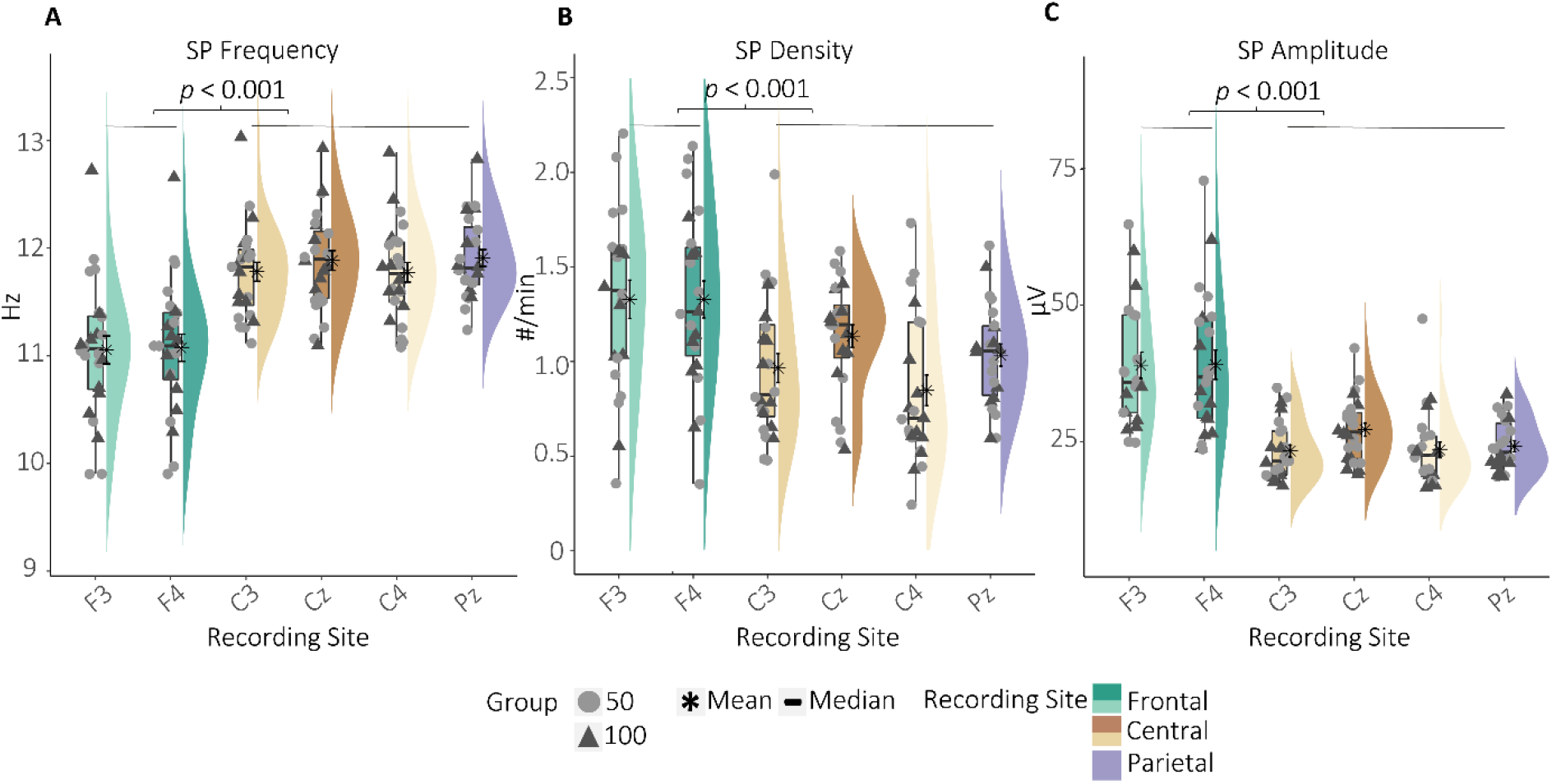
Individually identified SP (A) frequency, (B) density, and (C) amplitude at frontal, central, and parietal electrodes averaged across the two nights. *P-*values represent the results from Wilcoxon signed-rank tests comparing the average frontal and centro-parietal measures averaged across the two nights (Supplementary Table S4B). Frontal SPs differed from centro-parietal SPs in all three measures, implying the presence of a dominant slow frontal and a fast centro-parietal SP type.

The mean frequency of averaged slow frontal SPs was 11.07 Hz (min = 9.98 Hz, max = 12.67 Hz) while the averaged fast centro-parietal SPs had a mean frequency of 11.85 Hz (min = 11.15 Hz, max = 13.02 Hz). Note that even though (a) separate peaks were identifiable and (b) slow frontal and fast centro-parietal SPs differed reliably in their peak frequency, the fast centro-parietal SPs in our pre-school children were still below the typical fast SP frequency range in adults (Andrillon et al., 2011; Klinzing et al., 2016; Mölle et al., 2011).

Spearman’s rank correlations indicated that higher frequency, density, and amplitude of slow frontal SPs was associated with higher corresponding values of fast centro-parietal SPs (*ρ_frequency_* = 0.53, *p* = 0.008, *CI_2.5, 97.5_* [0.17, 0.76], *ρ_density_* = 0.70, *p* < 0.001, *CI_2.5, 97.5_* [0.37, 0.88], *ρ_amplitude_* = 0.81, *p* < 0.001, *CI_2.5, 97.5_* [0.57, 0.93]). Furthermore, neither the percentage of NREM sleep nor age were associated with slow frontal or fast centro-parietal SP frequency, density, or amplitude (−0.14 < *ρ* < 0.29, all *p* > 0.174, Supplementary Figure S4). In sum, individually identified SPs in frontal and centro-parietal sites differed in their characteristics, indicating that a slow frontal and a fast centro-parietal SP type is already present in 5- to 6-year-old children.

### 3.2 No Evidence for an Anterior or Posterior Predominance of Slow Oscillations

Since slow rhythmic neuronal activity during sleep shows a posterior rather than an anterior prevalence in children (Kurth, Ringli et al., 2010), we compared SO characteristics in averaged frontal (F3 & F4), averaged midline centro-parietal (Cz & Pz), and midline occipital (Oz) regions to examine any potential anterior or posterior predominance that might affect our following SP-SO coupling analyses. We decided to concentrate on midline derivations whenever possible, as SOs tend to travel along a midline anterior-posterior path (Murphy et al., 2009). We conducted separate repeated-measure ANOVAs on SO frequency, density, and amplitude with the within-person factors NIGHT (baseline vs. learning) and TOPOGRAPHY (frontal, centro-parietal, occipital).

SOs differed neither in their frequency (*F*(2,44) = 0.12, *p* = 0.888, *η^2^_G_*< 0.01), density (*F*(2,44) = 2.01, *p* = 0.147, *η^2^_G_* = 0.02), nor amplitude (*F*(2,44) = 2.14, *p* = 0.129, *η^2^_G_* = 0.03) among topographical locations (Figure 5). This effect did not differ between baseline and learning night (interaction effects: *F_frequency_*(2,44) = 0.65, *p* = 0.529, *η^2^_G_*< 0.01; *F_density_*(2,44) = 1.11, *p* = 0.337, *η^2^_G_*< 0.01; *F_amplitude_*(2,44) = 0.74, *p* = 0.482, *η^2^_G_* < 0.01). Furthermore, frequency and amplitude did not differ between the two PSG nights (*F_frequency_(*1,22) = 1.82, *p* = 0.191, *η^2^_G_*< 0.01; *F_amplitude_*(1,22) = 3.65, *p* = 0.069, *η^2^_G_* < 0.01). However, density was significantly higher during the baseline than the learning night (*F*(1,22) = 5.85, *p* = 0.024, *η^2^_G_*= 0.03). Taken together, our analyses did not indicate any topographical predominance of SOs in the present pre-school sample.

**Figure 5.**
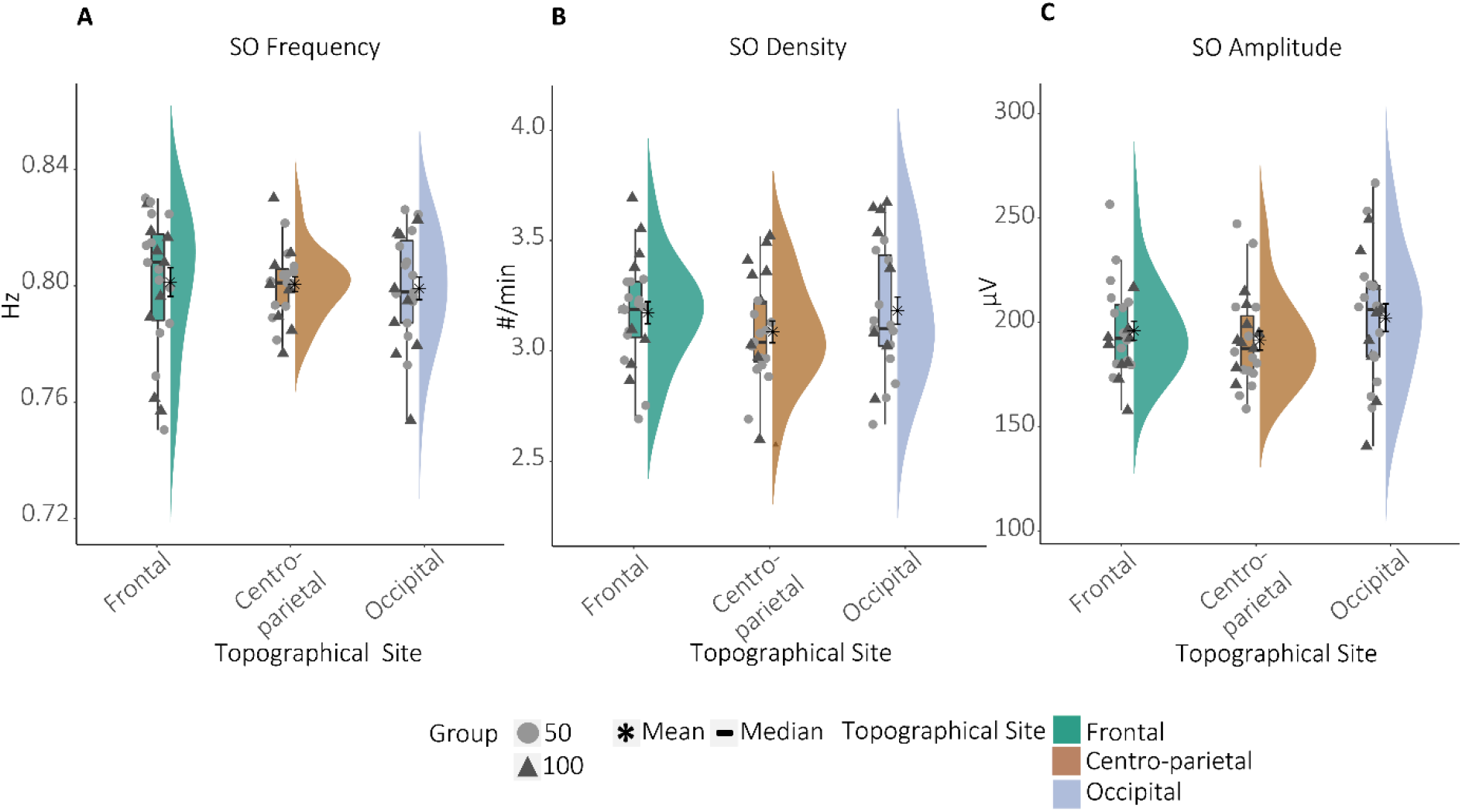
Main effects of the topographical differences for SO (A) frequency, (B) density, and (C) amplitude. Main effects from separate repeated-measure ANOVAs indicated no differences between topographical sites for any SO measure.

### 3.3 Exploration of Sleep Spindle Modulation During Slow Oscillations

#### 3.3.1 Power Modulations During Slow Oscillations

Having established the existence of separable slow frontal and fast centro-parietal SPs, we were interested in their temporal relation to SOs. In a first step, we explored SP power modulation during SOs on a descriptive level by contrasting power (5–20 Hz) during SOs (centred ± 1.2 s around the DOWN peak) with power during trials without SOs (Muehlroth et al., 2019). As there was no evidence for an anterior or posterior predominance of SOs, we examined frontal and centro-parietal SP power during averaged frontal (F3, F4), averaged midline centro-parietal (Cz, Pz), and occipital SOs (Oz). Cluster-based permutation tests revealed one cluster of increased power during SOs for both frontal and centro-parietal SP power (all cluster *p*s < 0.001). SP power was increased across the entire SP frequency range (9–16 Hz, Figure 6, dashed outline) and basically across the whole SO interval (Figure 6; for results for frontal and occipital SOs, see Supplementary Figure S5). Although this effect was apparent across SOs in all recording sites, it seemed most pronounced for centro-parietal SOs. The strongest frontal and centro-parietal SP power enhancements during SOs were observed during the transition from the DOWN to the subsequent UP peak in frequencies ≥12 Hz. Specifically for centro-parietal SOs, this enhanced power in the adult-like fast-SP range (12–15 Hz, Mölle et al., 2011; 13–16 Hz; Anderer et al., 2001; Schabus et al., 2007) was maintained throughout the UP state.

**Figure 6.**
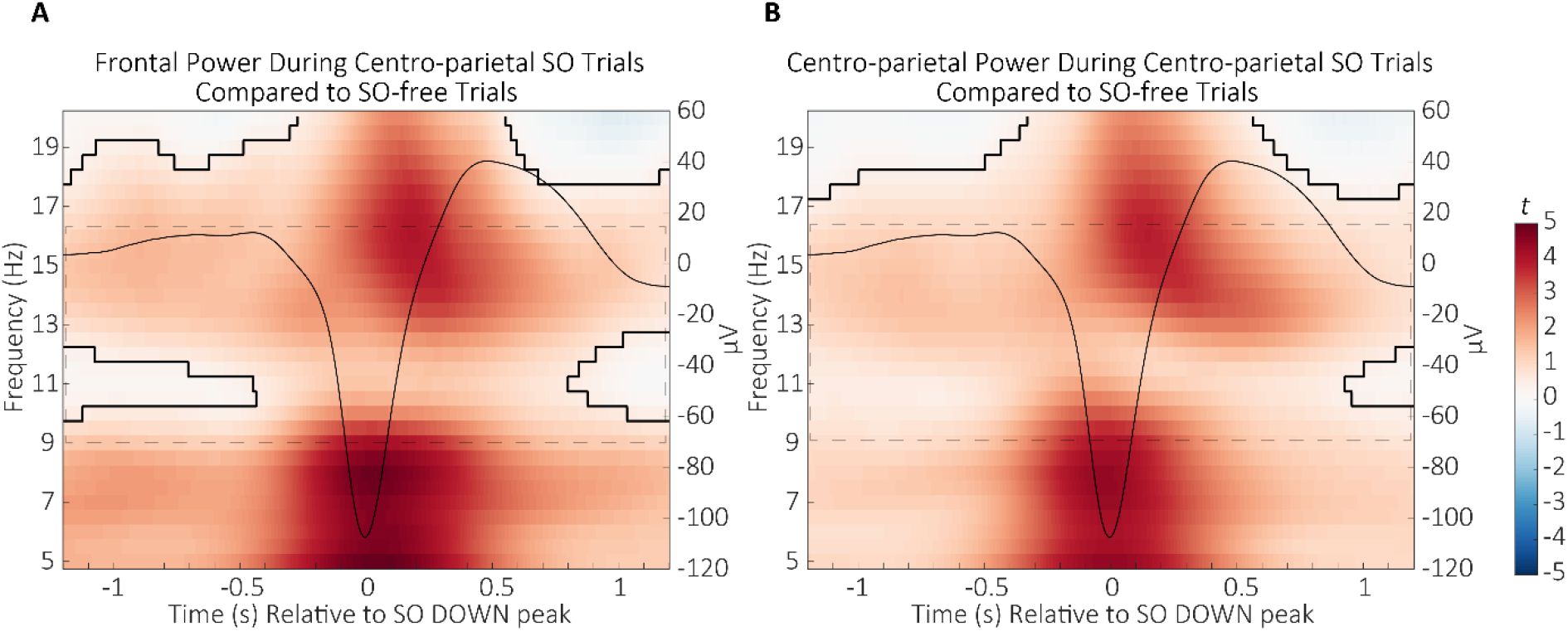
Differences in (A) frontal and (B) centro-parietal wavelet power during centro-parietal SOs compared to trials without SOs (*t*-score units). Significant clusters (cluster-based permutation test, cluster *α* < 0.05, two-sided test) are outlined. The SP frequency range is indicated by the reference window outlined in dashed lines. The average centro-parietal SO is projected onto the power differences to illustrate their relation to the SO phase (scale in µV on the right side of each time-frequency plot).

To summarise, we observed enhanced frontal and centro-parietal SP power during SOs, with a strong increase in power in the adult-like fast SP frequency range before and during the SO UP peak. Hence, we observed evidence for SP-SO coupling in pre-school children. On a descriptive level, the peak increase in the adult-like fast SP frequency range observed here appears slightly earlier than what is reported in the adult literature (Helfrich et al., 2018; Klinzing et al., 2016; Muehlroth et al., 2019).

#### 3.3.2 Modulation of Discrete Sleep Spindles During Slow Oscillations

Despite apparent power modulations within the SP frequency range during SOs, it is important to stress that these results do not necessarily reflect modulations of discrete individually identified slow frontal and fast centro-parietal SPs, given that for both SP types the mean frequency was identified at a lower frequency range (see SP results). In a second step, we were therefore interested in how the occurrence of individually identified SPs was related to the SO cycle.

To examine whether SPs and SOs actually co-occurred, we firstly determined the percentage of slow frontal and fast centro-parietal SP centres (SO DOWN peaks) occurring within an interval ± 1.2 s around the DOWN peak of SOs (SP centres, Muehlroth et al., 2019). We tested differences in SP-SO co-occurrence across SP types and SOs recorded in different locations using two separate repeated-measure ANOVAs with the within-person factors SP TYPE (slow frontal, fast centro-parietal) and SO TOPOGRAPHY (frontal, centro-parietal, occipital) on the percentage of SP events during SOs (SO DOWN peaks during SPs).

The percentage of SP centres co-occurring with SOs was overall significantly different between slow frontal and fast centro-parietal SPs (*F*(1,23) = 21.59, *p* < 0.001, *η^2^_G_*= 0.11) and across SOs in different topographical locations (*F*(1.49,34.35) = 103.25, *p* < 0.001, *η^2^_G_*= 0.32). Furthermore, the difference in SP centre occurrence between SP types was modulated by the topographical location of SOs (interaction effect: *F*(2,46) = 4.48, *p* = 0.017, *η^2^_G_*< 0.01). Post-hoc tests showed that the percentage of both slow frontal and fast centro-parietal SPs was considerably higher during SOs in centro-parietal compared to frontal and occipital recording sites (all *Z* < −3.00, all *p* < 0.001, Figure 7A; Supplementary Table S5A). Furthermore, in line with a general predominance of slow frontal SPs, a significantly higher percentage of slow frontal SPs, compared to fast centro-parietal SPs, co-occurred with frontal, centro-parietal, and occipital SOs (all *Z* < −3.00, all *p* < 0.001, Figure 7A, Supplementary Table S5B). Results for SOs co-occurring with SPs revealed similar results (Figure 7B, Supplementary Table S6).

**Figure 7.**
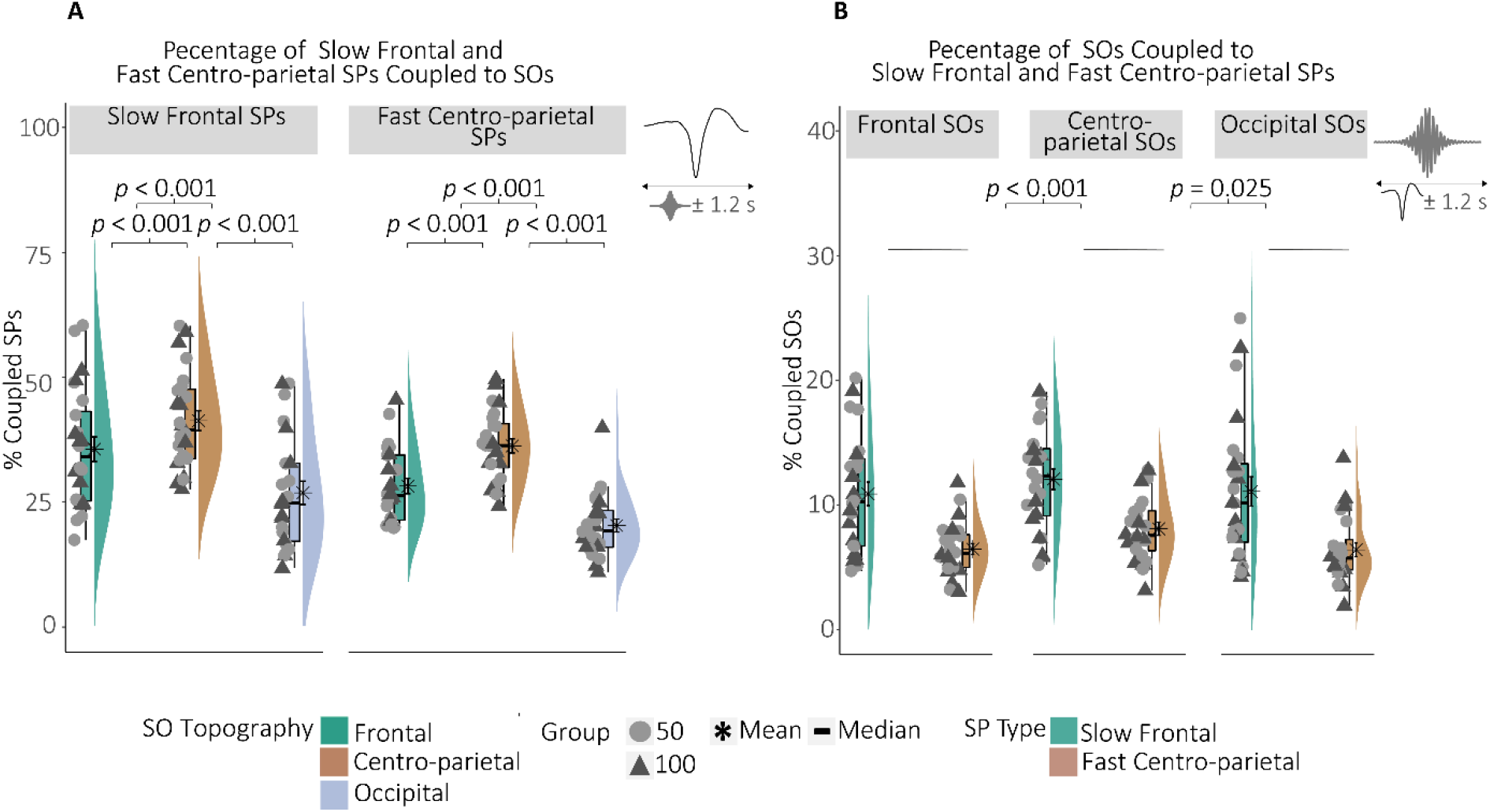
(A) Percentage of individually identified slow frontal and fast centro-parietal SPs co-occurring with frontal, centro-parietal, and occipital SOs. *P*-values represent the results from the post-hoc Wilcoxon signed-rank tests. (B) Percentage of frontal, centro-parietal, and occipital SOs co-occurring with individually identified slow frontal and fast centro-parietal SPs. *P*-values represent the results from the post-hoc Wilcoxon signed-rank tests on the main effect “SO Topography” comparing centro-parietal SO DOWN peak co-occurrence with individually identified SPs against frontal and occipital SO DOWN peak co-occurrence with individually identified SPs.

In sum, our analyses show that individually identified SPs generally co-occur with SOs, with more slow frontal SPs coinciding with SOs and centro-parietal SOs showing the highest coincidence with SPs.

Given the general presence of individually identified SPs during SOs, we were interested in the precise temporal modulation of these SPs during the SO cycle. Thus, we separately determined the percentage of slow frontal and fast centro-parietal SPs within specific 100 ms bins during an interval of ± 1.2 s around SO DOWN peaks to generate PETHs. To assess whether the modulation of SP occurrence within a bin was specific to the SO cycle, we compared the percentage distribution of SP centre occurrence with its randomly shuffled surrogate. Given the higher co-occurrence of SPs with centro-parietal SOs, we focus on results for centro-parietal SOs (for results for frontal and occipital SOs, see Supplementary Figure S6).

For fast centro-parietal SPs, we found an increased occurrence during the SO UP peak preceding the DOWN peak (cluster *p* = 0.002; −700 ms to −400 ms; UP peaks = −453.00 ms & 484.400 ms) and an attenuated incidence during the end of the SO (cluster *p* = 0.003, 900 ms to 1200 ms, Figure 8B). Similarly, slow frontal SPs were reduced during the end of the SO cycle (cluster *p* = 0.002, 1000 ms to 1200 ms, Figure 8A). However, the observed modulation of individually identified slow frontal and fast centro-parietal SPs during SOs does not look strong and matches neither the previous time-frequency results nor what we would expect from the adult literature (e.g., Muehlroth et al., 2019).

**Figure 8.**
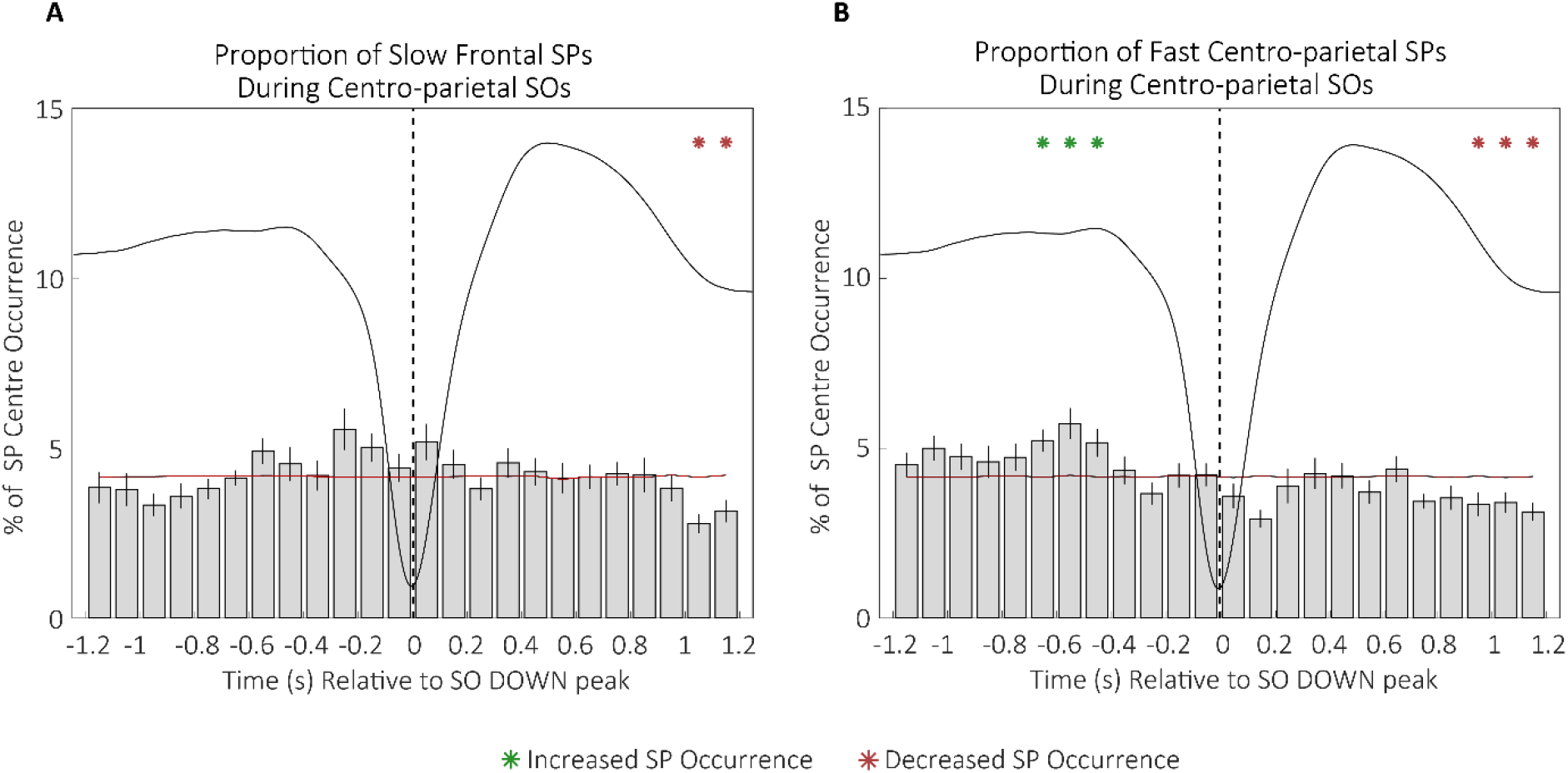
Percentage of individually identified (A) slow frontal and (B) fast centro-parietal SPs occurring within 100 ms bins during centro-parietal SOs. Green asterisks indicate increased SP occurrence (positive cluster, cluster *α* < 0.05, two-sided test) and red asterisks indicate decreased SP occurrence (negative cluster, cluster *α* < 0.05, two-sided test) compared to random occurrence (black horizontal line with standard error of the mean indicated in red). Error bars represent standard errors. The dashed vertical line represents the SO DOWN peak. The average centro-parietal SO is depicted in black.

Given that SP power modulation in the time-frequency analyses happened to be most pronounced in a range higher than the individually identified SPs, we exploratorily extracted SPs specifically higher in frequency than our individually defined upper SP frequency boundaries (“high” SPs) and repeated the PETH analyses. These discrete events should reflect the frequency range of peak power modulations in our time-frequency analyses more accurately. Indeed, “high” frontal SPs showed a mean frequency of 13.18 Hz (min = 12.31, max = 14.09 Hz) and “high” centro-parietal SPs had an average frequency of 13.58 Hz (min = 12.83, max = 14.27 Hz, see Supplementary Table S3 for further descriptive measures of “high” SPs). Even though we could not identify a dominant peak in the “high” SP frequency range in any of the participants’ power spectra, this does not prove the complete absence of such adult-like fast SPs (Supplementary Figure S3 for overall spectra; Supplementary Figure S7 for power spectra during trials with “high” SPs only). In particular, the results of the time-frequency analyses show that there might be a small number of discrete SPs in this range not powerful enough to elicit a peak in the power spectrum. However, like the fast SPs seen in adults, these SPs might still co-occur with SOs and be relevant to behaviour (see Supplementary Figure S8 and Supplementary Tables S7 and S8 for general co-occurrence results).

Cluster-based permutation tests did not reveal any statistically significant modulation of “high” frontal SP occurrence during centro-parietal SOs (Figure 9A; for results for frontal and occipital SOs, see Supplementary Figure S9). However, “high” centro-parietal SPs showed a pattern of increased SP occurrence before the UP peak preceding the DOWN peak (cluster *p* = 0.011; −1000 ms to −800 ms) and during the transition from the DOWN to the successive UP peak, including the UP peak (cluster *p* = 0.004, 100 ms to 400 ms). Furthermore, “high” centro-parietal SPs were attenuated starting before the UP peak prior to the DOWN peak lasting throughout the transition into the DOWN peak (cluster *p* < 0.001, −500 ms to −100 ms) and during the transition from the UP peak following the DOWN peak until the end of the SO cycle (cluster *p* < 0.001, 800 ms to 1200 ms, Figure 9B). This pattern reflects the power modulations of the time-frequency analyses more closely than the pattern of individually identified SPs does, supporting the notion of slightly earlier adult-like fast SP modulation in pre-school children during the SO cycle.

**Figure 9.**
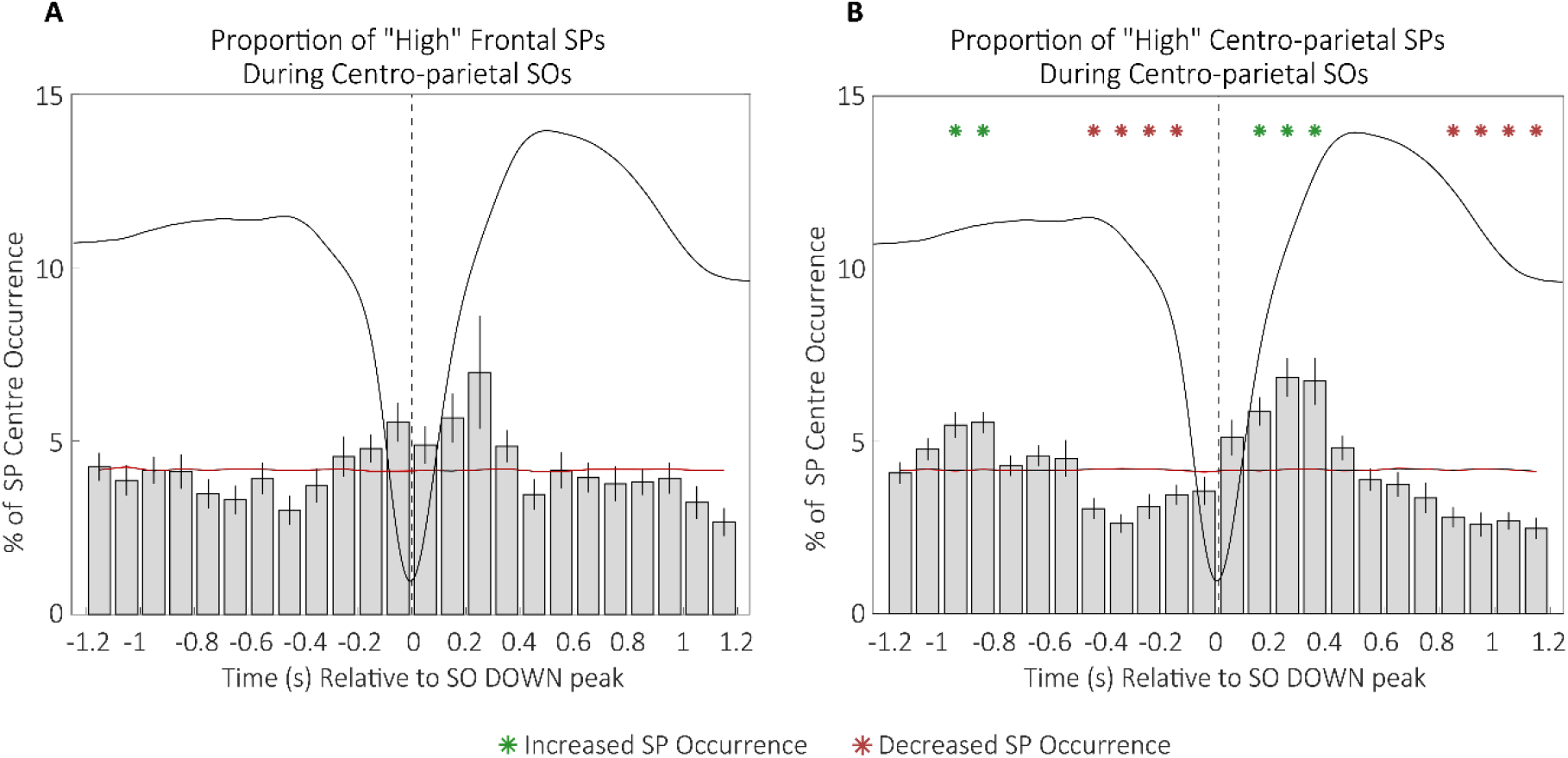
Percentage of “high” (A) frontal and (B) centro-parietal SPs occurring within 100 ms bins during centro-parietal SOs. Green asterisks indicate increased SP occurrence (positive cluster, cluster *α* < 0.05, two-sided test) and red asterisks indicate decreased SP occurrence (negative cluster, cluster *α* < 0.05, two-sided test) compared to random occurrence (black horizontal line with standard error of the mean indicated in red). Error bars represent standard errors. The dashed vertical line represents the SO DOWN peak. The average centro-parietal SO is depicted in black.

### 3.4 Recall Success After a Night of Sleep is Contingent on Memory Quality

Having established the presence of a slow frontal and a fast centro-parietal SP type and a modulation of SPs by SOs, we were interested in their relation with memory consolidation. As our associative memory task allows us to distinguish memories based on their learning trajectory during the evening into memories of varying quality, we firstly tested whether there was a difference in memory consolidation for low-, medium-, and high-quality memories. We applied a mixed factorial ANOVA with the between-person factor GROUP (Group_50_, Group_100_) and the within-person factor MEMORY QUALITY (low, medium, high) on the percentage of remembered items during morning recall. Similar to previous analyses on the difference between groups in general recall performance (Supplementary Figure S1, Supplementary Table S2), groups did not differ in the magnitude of memory consolidation (*F*(1,22) = 0.32, *p* = 0.580,*η^2^_G_* < 0.01). However, the extent of memory consolidation was different for low-, medium-, and high-quality memories (*F*(2,44) = 182.93, *p* < 0.001, *η^2^_G_*= 0.85). Overall, post-hoc tests revealed that the percentage of remembered items after a night of sleep was highest for items of high memory quality as compared to medium-(*Z* = −3.13, *p* = 0.002, *CI_2.5, 97.5_* [−3.91, −1.257]) and low-quality memories (*Z* = −4.24, *p* < 0.001, *CI_2.5, 97.5_* [−4.10, − 3.90]). Furthermore, successful recall of medium-quality memories was higher than that of low-quality memories (*Z* = −5.30, *p* < 0.001, *CI_2.5, 97.5_* [−4.10, −4.09], Figure 10A). The difference in the consolidation of memories of varying quality did not differ by group (interaction effect: *F*(2,44) = 0.27, *p* = 0.769, *η^2^_G_* < 0.01). Moreover, Spearman’s rank correlations revealed that the consolidation rates of low-, medium-, and high-quality memories were not correlated (*ρ_low*medium_* = 0.11, *p* = 0.618, *CI_2.5, 97.5_* [−0.35, 0.53], *ρ_low*high_* = −0.12, *p* = 0.575, *CI_2.5, 97.5_* [−0.54, 0.30], *ρ_medium*high_* = 0.18, *p*=0.400, *CI_2.5, 97.5_* [−0.19, 0.51]).

**Figure 10.**
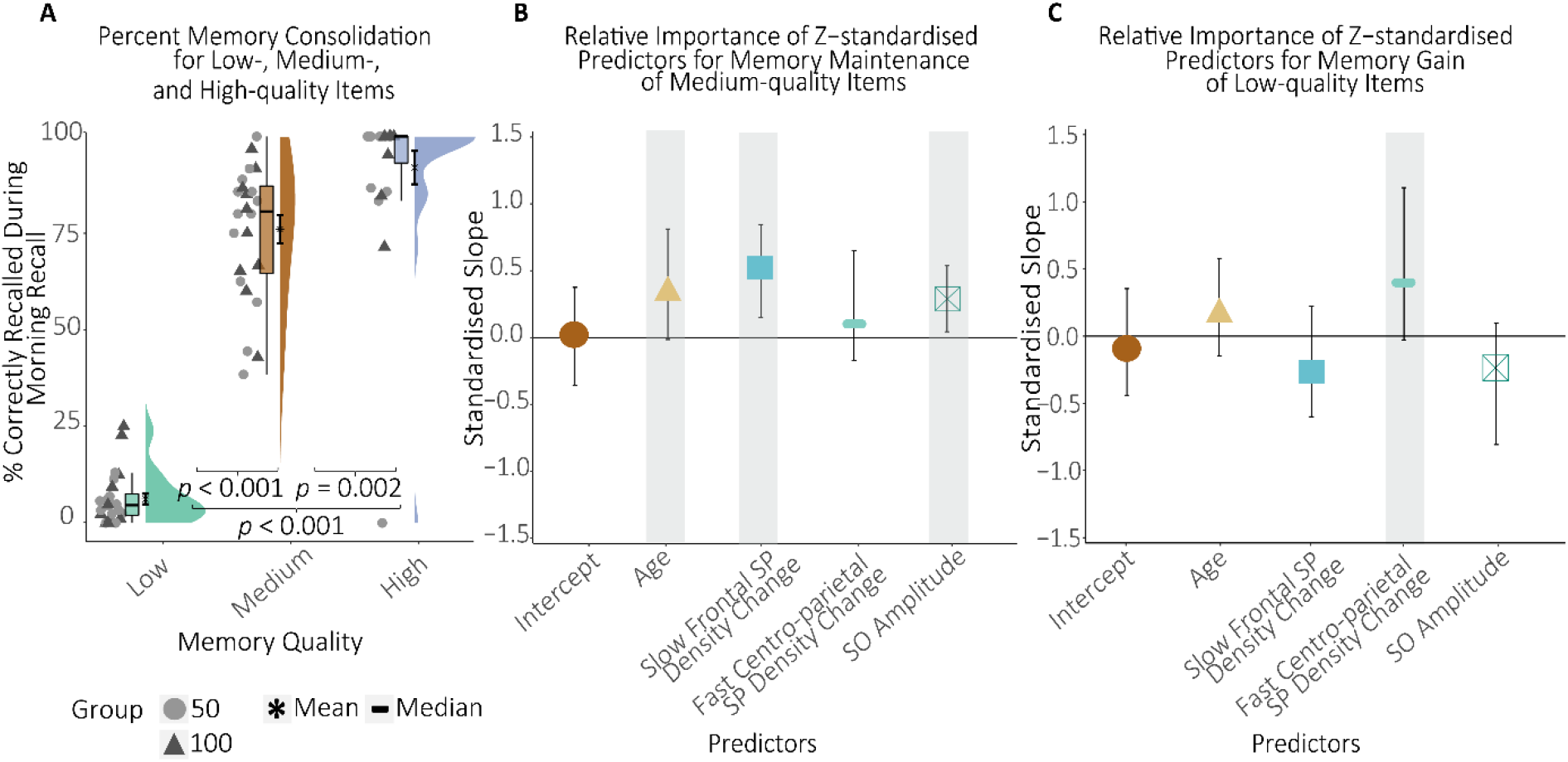
(A) Effect of sleep on memory consolidation for memories of varying quality. The better the memory quality, the more items were recalled after one night of sleep. *P-*values represent the results from post-hoc Wilcoxon signed-rank tests. (B & C) *Z*-standardised regression coefficients for memory consolidation of (B) medium- and (C) low-quality memories with their 95% simple bootstrap percentile confidence interval. Significant predictors are highlighted by the grey boxes. (B) Higher age, higher slow frontal SP density during the learning compared to the baseline night (Slow Frontal SP Density Change), and higher SO power were associated with stronger consolidation of medium-quality memories. (C) Higher fast centro-parietal SP density during the learning compared to the baseline night (Fast Centro-parietal SP Density Change) was associated with stronger memory consolidation of low-quality memories.

In sum, the extent of memory consolidation was contingent on memory quality with recall success after one night of sleep increasing with higher memory quality.

### 3.5 Individually Identified Slow Frontal and Fast Centro-parietal Sleep Spindles are Associated With Memory Consolidation

Since previous research in adults and children suggested that not only the mere presence but especially the learning-induced change in SP density is linked to the extent of sleep-associated memory consolidation (Friedrich et al., 2019; Gais et al., 2002; Lustenberger et al., 2015; Schabus et al., 2004), we calculated separate difference scores of SP density between the two PSG nights for individually identified slow frontal and fast centro-parietal SPs respectively. A positive difference score represents higher SP density during the learning as compared to the baseline night.

For slow rhythmic neuronal activity, power measures are usually associated with memory consolidation (Marshall et al., 2006). Therefore, we took the average SO amplitude across frontal and midline recording sites during the learning night as our measure of interest. We then examined the effect of slow frontal and fast centro-parietal SP density change and SO amplitude on the consolidation of low-, medium-, and high-quality memories using separate bootstrapped robust regressions. To control for a potential influence of chronological age on sleep-memory associations, age was included as a covariate. All variables were *Z*-standardised to enhance interpretability of bootstrap percentile CIs around the regression coefficients.

With respect to consolidation of medium-quality memories, results revealed that, besides age (*β* = 0.36, *p* = 0.020, *CI_2.5, 97.5_* [−0.02, 0.81]), higher slow frontal SP density change from baseline to learning night (*β* = 0.55, *p* < 0.001, *CI_2.5, 97.5_* [0.15, 0.85]) and SO amplitude (*β* = 0.29, *p* = 0.038, *CI_2.5, 97.5_* [0.04, 0.54]) were reliably associated with their higher maintenance (Figure 10B).

With regard to consolidation of low-quality memories, the increase in fast centro-parietal SP density during the learning night was significantly associated with a higher recall success (*β* = 0.39, *p* = 0.033, *CI_2.5, 97.5_* [−0.03, 1.10]), i.e., a memory gain (Figure 10C). We did not observe associations between electrophysiological sleep markers and maintenance of high-quality memories (a complete listing of all regression results can be found in Supplementary Tables S9–S11).

Taken together, age, learning-related slow frontal SP density change, and SO amplitude were related to memory maintenance of medium-quality memories while experience-related increase in fast centro-parietal SP density was associated with memory gain of low-quality memories.

### 3.6 Exploratory Analysis of the Association Between Sleep Spindle Modulation by Slow Oscillations With Memory Consolidation

The most prominent views on system consolidation suggest that the precise modulation of SPs during SOs, rather than their mere presence, represents a key mechanism underlying sleep-associated memory consolidation (Diekelmann & Born, 2010; Helfrich et al., 2018, 2019; Muehlroth et al., 2019). Hence, we asked whether and how the identified SP power modulations during SOs are associated with memory consolidation in the present sample of pre-schoolers. As the time-frequency analyses revealed a cluster of increased power during the whole SO cycle in a broad frequency range, we restricted our exploratory correlation analyses to the peak power increase during SOs, which covers the strong increase in the adult-like fast SP frequency range (Figure 6, Supplementary Figure S5). We therefore identified a mask of the 5% highest *t*-values in the range of 11–20 Hz (based on the increase visible in Figure 6) from the group-level contrast. These *t*-values were in a frequency range of 14–17.5 Hz. The mask was used to extract the *t*-values within this area for each participant. Afterwards the *Z*-standardised averaged value was correlated with memory consolidation of low-, medium-, and high-quality memories. These exploratory correlations did not yield any significant associations (Supplementary Figure S10A, all −0.3 < *ρ* < 0.3; all *p* > 0.178; see Supplementary Figure S10B and S10C for all exploratory results on an association between indicators of SP-SO coupling and memory consolidation).

## 4 Discussion

The present study aimed to characterise slow and fast SPs, their temporal interaction with SOs, and their relation to behavioural indicators of memory consolidation in pre-school children. Employing individualised rhythm detection methods, we found evidence for two separable SP types: A faster SP type in centro-parietal areas in addition to a more numerous, slower SP type in frontal sites. Individually identified fast centro-parietal SPs were nested in the adult-like slow SP range and already slightly modulated by the SO cycle. Surprisingly, we observed a clearer modulation of SPs higher than the individually identified SPs, roughly matching the adult-like fast SP range, during centro-parietal SOs. This modulation pattern seemed to be comparable to similar observations in adults, though with adult-like fast SPs in children peaking earlier than expected. While the pattern of SP modulation during SOs was not related to memory consolidation, importantly, SO amplitude and the increase in individually identified slow frontal SPs was reliably associated with sleep-associated maintenance of medium-quality memories. Further, individually identified fast centro-parietal SPs promoted the gain of low-quality memories. Together, our results indicate that, although the core mechanisms of sleep-associated system memory consolidation are not yet fully mature in pre-school children, subprocesses in their development-specific expression (i.e., slow frontal and fast centro-parietal SPs) already support sleep-associated memory consolidation in childhood.

### 4.1 Slow Frontal and Fast Centro-parietal Sleep Spindles in Pre-school Children

In the system consolidation framework, fast SPs are proposed to be a central mechanism for sleep-associated memory consolidation (Peyrache & Seibt, 2020; Rasch & Born, 2013). While existing evidence indicates the dominance of slow SPs, the reliable presence of an inherent fast SP type is still elusive in children (D’Atri et al., 2018; Hoedlmoser et al., 2014). Based on distinguishable individual peaks in frontal and cento-parietal spectra, we found that individually identified SPs differed in frequency, amplitude, and density at anterior and posterior recording sites in the majority of pre-school children. Thus, consistent with previous studies that relied on adult-derived frequency-based approaches (D’Atri et al., 2018; Hahn et al., 2019), our results support the presence of a dominant slow frontal and a fast centro-parietal SP type in pre-school children.

Importantly, the individually identified, fast centro-parietal SPs were within the range of adult-like slow SPs and thus differed from those identified in previous studies comparing slow and fast SPs in children (D’Atri et al., 2018; Hahn et al., 2019). However, the mean frequency of the individually identified fast centro-parietal SPs matches other studies that detected SPs individually in centro-parietal sites in 2- to 5-year-olds (Kurdziel et al., 2013; Olbrich et al., 2017). Further, it aligns well with findings demonstrating that SP frequency in general, and specifically in centro-parietal areas, is slower during childhood, increasing over the course of maturation (Campbell & Feinberg, 2009; Shinomiya et al., 1999). Hence, these observations may indicate that canonical fast SPs are not yet fully mature in pre-school children.

It should be noted that in most children, due to their proximity, two peaks were only identifiable upon separation of the power spectrum based on topography. Further, a small number of children expressed identical peak frequencies.

In general, varying features of neural rhythms likely reflect anatomical and functional properties of their underlying cortical and subcortical circuits (Andrillon et al., 2011; Buzsáki, 2006; Campbell & Feinberg, 2009; Piantoni, Poil et al., 2013; Saletin et al., 2013). Thus, two scenarios implying slightly different underlying developmental mechanisms are likely to account for developmental differences in the expression of fast centro-parietal SPs. Firstly, brain morphology undergoes strong developmental remodelling (Barnea-Goraly et al., 2005; Casey et al., 2000) and increasingly accelerated neuronal transmission allows for faster central processing. Hence, pruning (Campbell & Feinberg, 2009) and increased myelination (Nunez, 2000) in thalamo-cortical circuitries, and a decreasing degree of thalamic hyperpolarization (Andrillon et al., 2011; Steriade & Llinás, 1988) could directly account for frequency acceleration of fast SPs in the course of maturation. Secondly, it is also conceivable that changes in the generation mechanisms of SPs result in an increasing expression of SPs in the adult-like fast SP range across maturation. This would lead to power gains in the respective fast SP frequency range enabling the detectability of a peak once a sufficient number of fast SPs is expressed, and also leading to findings of increased frequency in the broad SP band. Thus, the individually identified fast centro-parietal SPs in pre-school children could either reflect a slower expression of adult-like fast SPs or a distinct rhythm that is no longer present in adults. However, disentangling the two possibilities necessitates longitudinal studies with combined electrophysiological and anatomical recordings (Lindenberger et al., 2011).

Taken together, we found evidence for two dissociable SP types in pre-school children. Given the nesting of fast centro-parietal SPs within the adult-like slow SP band, our results further support the utility of individualised approaches (Cox et al., 2017; Mölle et al., 2011; Ujma et al., 2015) to uncover true rhythmic neuronal activity in pre-school children.

### 4.2 Temporal Relation Between Sleep Spindles and Slow Oscillations

Recently it has been proposed that a precise modulation of SPs by the SO UP state might already be inherent to pre-pubertal children (aged around 8–-11 years; Hahn et al., 2020; Piantoni, Astill et al., 2013), though less pronounced and growing stronger across maturation (Hahn et al., 2020). Indeed, extending these results to pre-school children, we did observe a slight modulation of fast centro-parietal SP occurrence by the SO UP state that was not apparent for slow frontal SPs. Surprisingly, the power and occurrence of centro-parietal SPs in frequencies higher than the individually identified SPs, matching the adult-like fast SP range, exhibited an even more pronounced modulation during SOs. This seems to be comparable to the SP-SO coupling identified in adults, albeit occurring slightly earlier (Klinzing et al., 2016; Muehlroth et al., 2019). The less precise co-occurrence of “high” SPs with the UP state of SOs fits findings of SP-SO dispersion during aging (Helfrich et al., 2018; Muehlroth et al., 2019) and suggests that together with the number of adult-like fast SPs, their precise timing in relation to the SO UP peak still needs to mature in childhood development.

Hence, the patterns of individually identified fast centro-parietal SP coupling and of “high” centro-parietal SP modulation seem to imply that two distinct mechanisms form the basis for development of strong, precise SP-SO coupling (i.e., growing strength as well as increasing temporal precision). While we cannot answer which of these mechanisms leads to the fully mature SP-SO coupling seen in adults, our results cautiously suggest that they might act in concert. After all, it appears that the greater presence of adult-like fast SPs renders SP-SO coupling more precise and pronounced. It remains unclear which of the involved components, SPs, SOs, or both underlie developmental differences in SP-SO coupling.

Besides the maturation of adult-like fast SPs, the development of SOs themselves could contribute to the above findings. SOs are more numerous and powerful during younger age and do not yet show a prefrontal dominance (Kurth, Jenni et al., 2010, Kurth, Ringli et al., 2010). While we neither observed a frontal nor the expected central or posterior dominance of SOs (Kurth, Ringli et al., 2010; Timofeev et al., 2020), we found more SPs and a clearer SP modulation pattern for centro-parietal SOs. In previous work, however, it was precisely the co-ordination of fast SPs with frontal SOs (Hahn et al., 2020; Helfrich et al., 2018, 2019; Muehlroth et al., 2019) and, in children, frontal SOs in general (Prehn-Kristensen et al., 2014) that were found to be behaviourally relevant. One can only speculate that the relative lack of prefrontal SO dominance might affect the SP-SO modulation pattern in children. Further, since existing literature suggests that SOs are more powerful at younger ages (Campbell & Feinberg, 2009; Kurth, Jenni et al., 2010), it is possible that the level of depolarisation reached during the transition from the DOWN peak to the UP peak might suffice to elicit SPs in young children. This might account for the rise in “high” centro-parietal SPs already occurring before the UP peak. To sum up, we found evidence for a weak and imprecise modulation of fast centro-parietal SPs by SOs in pre-school children implying that overall, the hallmark system consolidation mechanism is not yet fully mature at this age.

### 4.3 Behavioural Relevance of Slow Frontal and Fast Centro-parietal Sleep Spindles and Their Pattern of Co-occurrence With Slow Oscillations for Memory Consolidation

The prevailing view proposes that the precise coupling of hippocampal ripples to fast SPs during the SO UP state supports system consolidation during sleep by providing a time window of enhanced cortical excitability that invokes the stabilisation of hippocampal mnemonic patterns in respective cortical areas (Clemens et al., 2007; Diekelmann & Born, 2010; Helfrich et al., 2019; Staresina et al., 2015). Despite observing a modulation of SPs during SOs in the current study, we did not find evidence of its critical contribution to memory consolidation in pre-school children. This may suggest that the coordinated triad of hippocampal ripples, SPs, and SOs is not only not fully developed, but also not yet behaviourally relevant in pre-school children. Overall, these observations suggest that over the course of development, neuronal mechanisms supporting sleep-associated system consolidation might require refined synchronisation to fully support stabilisation and integration of novel mnemonic contents.

While the precise interaction between SOs and SPs was not associated with memory consolidation, a greater increase in individually identified slow frontal and fast centro-parietal SPs during the learning night was related to stronger memory consolidation in pre-school children, independently of their co-occurrence with SOs. This resonates with findings demonstrating that both coupled and uncoupled SPs benefit memory in adults while the number of coupled SPs possibly needs to exceed a certain threshold to further ameliorate memory (Denis et al., 2020). Thus, it is likely that isolated SPs compensate for the lack and imprecision of SP-SO coupling, challenging the view of SP-SO coupling as the central mechanism of sleep-associated memory consolidation (Diekelmann & Born, 2010; Latchoumane et al., 2017). While our results do not rule out that more precise coordination of SPs and SOs provides additional advantages for memory consolidation, all in all, our results imply that slow frontal and fast centro-parietal SPs in their development-specific expression support memory consolidation independent of their modulation by SOs. Note that the order of baseline and experimental nights was not counterbalanced, so we cannot exclude the possibility that differences in SP density between the baseline and learning night were due to habituation effects or other non-learning related, non-systematic causes. However, sleep architecture did not differ between the two nights. Furthermore, we did not examine the effect of a wake retention interval on task performance. Hence, it cannot be excluded that different and/or additional mechanisms may also act on formed memories during wakefulness and that these could result in comparable behavioural effects as those observed across a retention interval of sleep.

### 4.4 Slow Frontal and Fast Centro-parietal Sleep Spindles are Differentially Associated With Memory Maintenance and Gain – Implications for Differential Functions?

Individually identified slow frontal and fast centro-parietal SPs were not only both linked to memory consolidation but showed a differential association with memory maintenance and gain. A stronger increase in slow frontal SPs during the learning night was reliably related to higher maintenance of medium-quality memories while the rise in individually identified fast centro-parietal SPs was associated with gains of low-quality memories. Importantly, a given memory representation needs to be accompanied by a certain level of hippocampal and cortical activation for system consolidation mechanisms to work (Schoch et al., 2017; Tucker & Fishbein, 2008). The lack of accessibility of low-quality items in the evening session does not necessarily indicate the absence of a mnemonic representation but might be due to retrieval-rooted factors such as impaired retrieval search (Ackerman, 1985), retrieval-induced forgetting (Aslan & Bäuml, 2010), and/or reduced attentional guidance. Thus, the gain effect for low-quality items most likely reflects the release from recall perturbing factors overnight rather than a sleep-associated emergence of novel memory representations (Fenn & Hambrick, 2013; Muehlroth et al., 2020; Nettersheim et al., 2015). Previously, weakly encoded items were linked to increased hippocampal reactivation and SPs during sleep, thereby indicating preferential system consolidation of those memories most prone to be forgotten (Denis et al., 2020; Schapiro et al., 2018). Hence, the increased availability of low-quality items after sleep could reflect a strengthening of a weak mnemonic trace and/or relief from retrieval inhibiting factors. Thus, fast centro-parietal SPs are not only functionally relevant, but may already be specifically involved in hippocampal-cortical system consolidation in pre-school children – even without top-down co-ordination by SOs.

While the role of slow SPs in memory consolidation is still open, they could very well also represent hippocampal-cortical integration that compensates for the absence of fast SP-SO coupling in children. Considering the data presented in this article, we cannot provide a conclusive explanation of the differential functions of fast and slow SPs for memory maintenance and gain and further studies are definitely required. However, it has been suggested that slow SPs might be involved in cortico-cortical rather than hippocampal-cortical communication (Astori et al., 2013; Doran, 2003; Rasch & Born, 2013; Timofeev & Chauvette, 2013). As the effect of sleep on memory consolidation depends on the level of cortical integration of a memory during encoding (Himmer et al., 2017), one could cautiously speculate that medium-quality items have already established a stronger cortical trace than low-quality memories. Hence, there might be less need for hippocampal-cortical communication than for cortico-cortical distribution for these memories, potentially reflecting a stabilisation process. The exact prerequisites and mechanisms for overnight gain and maintenance certainly warrant further interrogation. Nevertheless, the present results provide further evidence that SP-related processes contribute to overnight system-level consolidation, even in pre-school children.

### 4.5 Conclusions

Overall, the present results underscore the functional relevance of inherent slow frontal and fast centro-parietal SPs for memory consolidation in pre-school children, despite not fully developed SP-SO coupling. Notably, the development-specific expression of fast centro-parietal SPs was associated with sleep-associated memory gain while slow frontal SPs were related to memory maintenance.

## References

Ackerman, B. P. (1985). Constraints on retrieval search for episodic information in children and adults. Journal of Experimental Child Psychology, 40(1), 152–180. https://doi.org/10.1016/0022-0965(85)90070-0

Adamczyk, M. (2015). Automatic sleep spindle detection and genetic influence estimation using continuous wavelet transform. Frontiers in Human Neuroscience, 9, Article 20. https://doi.org/10.3389/fnhum.2015.00624

Anderer, P., Klösch, G., Gruber, G., Trenker, E., Pascual-Marqui, R. D., Zeitlhofer, J., Barbanoj, M. J., Rappelsberger, P., & Saletu, B. (2001). Low-resolution brain electromagnetic tomography revealed simultaneously active frontal and parietal sleep spindle sources in the human cortex. Neuroscience, 103(3), 581–592. https://doi.org/10.1016/S0306-4522(01)00028-8

Andrade, K. C., Spoormaker, V. I., Dresler, M., Wehrle, R., Holsboer, F., Samann, P. G., & Czisch, M. (2011). Sleep spindles and hippocampal functional connectivity in human NREM sleep. Journal of Neuroscience, 31(28), 10331–10339. https://doi.org/10.1523/JNEUROSCI.5660-10.2011

Andrillon, T., Nir, Y., Staba, R. J., Ferrarelli, F., Cirelli, C., Tononi, G., & Fried, I. (2011). Sleep spindles in humans: Insights from intracranial EEG and unit recordings. Journal of Neuroscience, 31(49), 17821–17834. https://doi.org/10.1523/JNEUROSCI.2604-11.2011

Aru, J., Aru, J., Priesemann, V., Wibral, M., Lana, L., Pipa, G., Singer, W., & Vicente, R. (2015). Untangling cross-frequency coupling in neuroscience. Current Opinion in Neurobiology, 31, 51–61. https://doi.org/10.1016/j.conb.2014.08.002

Ashworth, A., Hill, C. M., Karmiloff-Smith, A., & Dimitriou, D. (2014). Sleep enhances memory consolidation in children. Journal of Sleep Research, 23(3), 304–310. https://doi.org/10.1111/jsr.12119

Aslan, A., & Bäuml, K.-H. T. (2010). Retrieval-induced forgetting in young children. Psychonomic Bulletin & Review, 17(5), 704–709. https://doi.org/10.3758/PBR.17.5.704

Astori, S., Wimmer, R. D., & Lüthi, A. (2013). Manipulating sleep spindles: Expanding views on sleep, memory, and disease. Trends in Neurosciences, 36(12), 738–748. https://doi.org/10.1016/j.tins.2013.10.001

Ayoub, A., Aumann, D., Hörschelmann, A., Kouchekmanesch, A., Paul, P., Born, J., & Marshall, L. (2013). Differential effects on fast and slow spindle activity, and the sleep slow oscillation in humans with carbamazepine and flunarizine to antagonize voltage-dependent Na+ and Ca2+ channel activity. Sleep, 36(6), 905–911. https://doi.org/10.5665/sleep.2722

Backhaus, J., Hoeckesfeld, R., Born, J., Hohagen, F., & Junghanns, K. (2008). Immediate as well as delayed post learning sleep but not wakefulness enhances declarative memory consolidation in children. Neurobiology of Learning and Memory, 89(1), 76–80. https://doi.org/10.1016/j.nlm.2007.08.010

Barnea-Goraly, N., Menon, V., Eckert, M., Tamm, L., Bammer, R., Karchemskiy, A., Dant, C. C., & Reiss, A. L. (2005). White matter development during childhood and adolescence: A cross-sectional diffusion tensor imaging study. Cerebral Cortex, 15(12), 1848–1854. https://doi.org/10.1093/cercor/bhi062

Berry, R. B., Brooks, R., Harding, S. M., Marcus, C., & Vaugh, B. V. (2015). AASM Scoring Manual Version 2.2. 7. American Academy of Sleep Medicine.

Bonjean, M., Baker, T., Lemieux, M., Timofeev, I., Sejnowski, T., & Bazhenov, M. (2011). Corticothalamic feedback controls sleep spindle duration in vivo. Journal of Neuroscience, 31(25), 9124–9134. https://doi.org/10.1523/JNEUROSCI.0077-11.2011

Buzsáki, G. (1998). Memory consolidation during sleep: A neurophysiological perspective. Journal of Sleep Research, 7(S1), 17–23. https://doi.org/10.1046/j.1365-2869.7.s1.3.x

Buzsáki, G. (2006). Rhythms of the brain. Oxford University Press. https://doi.org/10.1093/acprof:oso/9780195301069.001.0001

Buzsáki, G., & Mizuseki, K. (2014). The log-dynamic brain: How skewed distributions affect network operations. Nature Reviews Neuroscience, 15(4), 264–278. https://doi.org/10.1038/nrn3687

Campbell, I. G., & Feinberg, I. (2009). Longitudinal trajectories of non-rapid eye movement delta and theta EEG as indicators of adolescent brain maturation. Proceedings of the National Academy of Sciences of the United States of America, 106(13), 5177–5180. https://doi.org/10.1073/pnas.0812947106

Campbell, I. G., & Feinberg, I. (2016). Maturational patterns of sigma frequency power across childhood and adolescence: A longitudinal study. Sleep, 39(1), 193–201. https://doi.org/10.5665/sleep.5346

Casey, B. J., Giedd, J. N., & Thomas, K. M. (2000). Structural and functional brain development and its relation to cognitive development. Biological Psychology, 54(1–3), 241–257. https://doi.org/10.1016/S0301-0511(00)00058-2

Clarke, A. R., Barry, R. J., McCarthy, R., & Selikowitz, M. (2001). Age and sex effects in the EEG: Development of the normal child. Clinical Neurophysiology, 112(5), 806–814. https://doi.org/10.1016/S1388-2457(01)00488-6

Clemens, Z., Molle, M., Eross, L., Barsi, P., Halasz, P., & Born, J. (2007). Temporal coupling of parahippocampal ripples, sleep spindles and slow oscillations in humans. Brain, 130(11), 2868–2878. https://doi.org/10.1093/brain/awm146

Contreras, D., Destexhe, A., Sejnowski, T. J., & Steriade, M. (1997). Spatiotemporal patterns of spindle oscillations in cortex and thalamus. The Journal of Neuroscience, 17(3), 1179–1196. https://doi.org/10.1523/JNEUROSCI.17-03-01179.1997

Cox, R., Schapiro, A. C., Manoach, D. S., & Stickgold, R. (2017). Individual differences in frequency and topography of slow and fast sleep spindles. Frontiers in Human Neuroscience, 11, Article 433. https://doi.org/10.3389/fnhum.2017.00433

Craik, F. I. M., & Lockhart, R. S. (1972). Levels of processing: A framework of memory research. Journal of Verbal Learning and Verbal Behavior, 11(6), 671–684. https://doi.org/10.1016/S0022-5371(72)80001-X

Danner, F. W., & Taylor, A. M. (1973). Integrated pictures and relational imagery training in children’s learning. Journal of Experimental Child Psychology, 16(1), 47–54. https://doi.org/10.1016/0022-0965(73)90061-1

D’Atri, A., Novelli, L., Ferrara, M., Bruni, O., & De Gennaro, L. (2018). Different maturational changes of fast and slow sleep spindles in the first four years of life. Sleep Medicine, 42, 73–82. https://doi.org/10.1016/j.sleep.2017.11.1138

De Gennaro, L., & Ferrara, M. (2003). Sleep spindles: An overview. Sleep Medicine Reviews, 7(5), 423– 440. https://doi.org/10.1053/smrv.2002.0252

Denis, D., Mylonas, D., Poskanzer, C., Bursal, V., Payne, J. D., & Stickgold, R. (2020). Sleep spindles facilitate selective memory consolidation. BioRxiv. https://doi.org/10.1101/2020.04.03.022434

Diekelmann, S., & Born, J. (2010). The memory function of sleep. Nature Reviews Neuroscience, 11(2), 114–126. https://doi.org/10.1038/nrn2762

Doran, S. M. (2003). The dynamic topography of individual sleep spindles. Sleep Research Online, 5(4), 133–139.

Drosopoulos, S., Schulze, C., Fischer, S., & Born, J. (2007). Sleep’s function in the spontaneous recovery and consolidation of memories. Journal of Experimental Psychology: General, 136(2), 169– 183. https://doi.org/10.1037/0096-3445.136.2.169

Dumay, N. (2016). Sleep not just protects memories against forgetting, it also makes them more accessible. Cortex, 74, 289–296. https://doi.org/10.1016/j.cortex.2015.06.007

Dumay, N. (2018). Look more carefully: Even your data show sleep makes memories more accessible. A reply to Schreiner and Rasch (2018). Cortex, 101, 288–293. https://doi.org/10.1016/j.cortex.2017.12.013

Eichstaedt, K. E., Kovatch, K., & Maroof, D. A. (2013). A less conservative method to adjust for familywise error rate in neuropsychological research: The Holm’s sequential Bonferroni procedure. NeuroRehabilitation, 32(3), 693–696. https://doi.org/10.3233/NRE-130893

Fandakova, Y., Sander, M. C., Grandy, T. H., Cabeza, R., Werkle-Bergner, M., & Shing, Y. L. (2018). Age differences in false memory: The importance of retrieval monitoring processes and their modulation by memory quality. Psychology and Aging, 33(1), 119–133. https://doi.org/10.1037/pag0000212

Fenn, K. M., & Hambrick, D. Z. (2013). What drives sleep-dependent memory consolidation: Greater gain or less loss? Psychonomic Bulletin & Review, 20(3), 501–506. https://doi.org/10.3758/s13423-012-0366-z

Fernandez, L. M. J., & Lüthi, A. (2020). Sleep spindles: Mechanisms and functions. Physiological Reviews, 100(2), 805–868. https://doi.org/10.1152/physrev.00042.2018

Friedrich, M., Mölle, M., Friederici, A. D., & Born, J. (2019). The reciprocal relation between sleep and memory in infancy: Memory-dependent adjustment of sleep spindles and spindle-dependent improvement of memories. Developmental Science, 22(2), Article e12743. https://doi.org/10.1111/desc.12743

Friedrich, M., Mölle, M., Friederici, A. D., & Born, J. (2020). Sleep-dependent memory consolidation in infants protects new episodic memories from existing semantic memories. Nature Communications, 11(1), Article 1298. https://doi.org/10.1038/s41467-020-14850-8

Friedrich, M., Wilhelm, I., Born, J., & Friederici, A. D. (2015). Generalization of word meanings during infant sleep. Nature Communications, 6(1), Article 6004. https://doi.org/10.1038/ncomms7004

Gais, S., Mölle, M., Helms, K., & Born, J. (2002). Learning-dependent increases in sleep spindle density. Journal of Neuroscience, 22(15), 6830–6834. https://doi.org/10.1523/JNEUROSCI.22-15-06830.2002

Goodman, R. (1997). The Strengths and Difficulties Questionnaire: A research note. Journal of Child Psychology and Psychiatry, 38(5), 581–586. https://doi.org/10.1111/j.1469-7610.1997.tb01545.x

Grandy, T. H., Werkle-Bergner, M., Chicherio, C., Schmiedek, F., Lövdén, M., & Lindenberger, U. (2013). Peak individual alpha frequency qualifies as a stable neurophysiological trait marker in healthy younger and older adults: Alpha stability. Psychophysiology, 50(6), 570–582. https://doi.org/10.1111/psyp.12043

Habib, R., & Nyberg, L. (2008). Neural correlates of availability and accessibility in memory. Cerebral Cortex, 18(7), 1720–1726. https://doi.org/10.1093/cercor/bhm201

Hahn, M., Heib, D., Schabus, M., Hoedlmoser, K., & Helfrich, R. F. (2020). Slow oscillation-spindle coupling predicts enhanced memory formation from childhood to adolescence. eLife, 9, Article e53730. https://doi.org/10.7554/eLife.53730

Hahn, M., Joechner, A.-K., Roell, J., Schabus, M., Heib, D. P., Gruber, G., Peigneux, P., & Hoedlmoser, K. (2019). Developmental changes of sleep spindels and their impact on sleep-dependent memory consolidation and general cognitive abilities: A longitudinal approach. Developmental Science, 22(1), Article e12706. https://doi.org/10.1111/desc.12706

Helfrich, R. F., Lendner, J. D., Mander, B. A., Guillen, H., Paff, M., Mnatsakanyan, L., Vadera, S., Walker, M. P., Lin, J. J., & Knight, R. T. (2019). Bidirectional prefrontal-hippocampal dynamics organize information transfer during sleep in humans. Nature Communications, 10(1), 3572. https://doi.org/10.1038/s41467-019-11444-x

Helfrich, R. F., Mander, B. A., Jagust, W. J., Knight, R. T., & Walker, M. P. (2018). Old bBrains come uncoupled in sleep: Slow wave-spindle synchrony, brain atrophy, and forgetting. Neuron, 97(1), 221–230.e4. https://doi.org/10.1016/j.neuron.2017.11.020

Himmer, L., Müller, E., Gais, S., & Schönauer, M. (2017). Sleep-mediated memory consolidation depends on the level of integration at encoding. Neurobiology of Learning and Memory, 137, 101–106. https://doi.org/10.1016/j.nlm.2016.11.019

Hoedlmoser, K., Heib, D. P. J., Roell, J., Peigneux, P., Sadeh, A., Gruber, G., & Schabus, M. (2014). Slow sleep spindle activity, declarative memory, and general cognitive abilities in children. Sleep, 37(9), 1501–1512. https://doi.org/10.5665/sleep.4000

Jankel, W. R., & Niedermeyer, E. (1985). Sleep spindles. Journal of Clinical Neurophysiology, 2(1), 1–35. https://doi.org/10.1097/00004691-198501000-00001

Jasper, H. H. (1958). The ten-twenty electrode system of the International Federation. Electroencephalography and Clinical Neurophysiology, 10, 370–375.

Kleiner, M., Brainard, D., & Pelli, D. (2007). What’s new in Psychtoolbox-3? 89. Perception 36 ECVP Abstract Supplement

Klinzing, J. G., Mölle, M., Weber, F., Supp, G., Hipp, J. F., Engel, A. K., & Born, J. (2016). Spindle activity phase-locked to sleep slow oscillations. NeuroImage, 134, 607–616. https://doi.org/10.1016/j.neuroimage.2016.04.031

Kosciessa, J. Q., Grandy, T. H., Garrett, D. D., & Werkle-Bergner, M. (2020). Single-trial characterization of neural rhythms: Potential and challenges. NeuroImage, 206, Article 116331. https://doi.org/10.1016/j.neuroimage.2019.116331

Kurdziel, L., Duclos, K., & Spencer, R. M. C. (2013). Sleep spindles in midday naps enhance learning in preschool children. Proceedings of the National Academy of Sciences of the United States of America, 110(43), 17267–17272. https://doi.org/10.1073/pnas.1306418110

Kurth, S., Jenni, O. G., Riedner, B. A., Tononi, G., Carskadon, M. A., & Huber, R. (2010). Characteristics of sleep slow waves in children and adolescents. Sleep, 33(4), 475–480. https://doi.org/10.1093/sleep/33.4.475

Kurth, S., Ringli, M., Geiger, A., LeBourgeois, M., Jenni, O. G., & Huber, R. (2010). Mapping of cortical activity in the first two decades of life: A high-density sleep electroencephalogram study. Journal of Neuroscience, 30(40), 13211–13219. https://doi.org/10.1523/JNEUROSCI.2532-10.2010

Latchoumane, C.-F. V., Ngo, H.-V. V., Born, J., & Shin, H.-S. (2017). Thalamic spindles promote memory formation during sleep through triple phase-locking of cortical, thalamic, and hippocampal rhythms. Neuron, 95(2), 424–435.e6. https://doi.org/10.1016/j.neuron.2017.06.025

Lindenberger, U., von Oertzen, T., Ghisletta, P., & Hertzog, C. (2011). Cross-sectional age variance extraction: What’s change got to do with it? Psychology and Aging, 26(1), 34–47. https://doi.org/10.1037/a0020525

Lustenberger, C., Wehrle, F., Tüshaus, L., Achermann, P., & Huber, R. (2015). The multidimensional aspects of sleep spindles and their relationship to word-pair memory consolidation. Sleep, 38(7), 1093–1103. https://doi.org/10.5665/sleep.4820

Lüthi, A. (2014). Sleep spindles: Where they come from, what they do. The Neuroscientist, 20(3), 243– 256. https://doi.org/10.1177/1073858413500854

Maris, E., & Oostenveld, R. (2007). Nonparametric statistical testing of EEG- and MEG-data. Journal of Neuroscience Methods, 164(1), 177–190. https://doi.org/10.1016/j.jneumeth.2007.03.024

Marshall, L., Helgadóttir, H., Mölle, M., & Born, J. (2006). Boosting slow oscillations during sleep potentiates memory. Nature, 444(7119), 610–613. https://doi.org/10.1038/nature05278

Marshall, P. J., Bar-Haim, Y., & Fox, N. A. (2002). Development of the EEG from 5 months to 4 years of age. Clinical Neurophysiology, 113(8), 1199–1208. https://doi.org/10.1016/S1388-2457(02)00163-3

Maski, K., Holbrook, H., Manoach, D., Hanson, E., Kapur, K., & Stickgold, R. (2015). Sleep dependent memory consolidation in children with autism spectrum disorder. Sleep, 38(12), 1955–1963. https://doi.org/10.5665/sleep.5248

Mölle, M., Bergmann, T. O., Marshall, L., & Born, J. (2011). Fast and slow spindles during the sleep slow oscillation: Disparate coalescence and engagement in memory Processing. Sleep, 34(10), 1411–1421. https://doi.org/10.5665/SLEEP.1290

Mölle, M., Eschenko, O., Gais, S., Sara, S. J., & Born, J. (2009). The influence of learning on sleep slow oscillations and associated spindles and ripples in humans and rats. European Journal of Neuroscience, 29(5), 1071–1081. https://doi.org/10.1111/j.1460-9568.2009.06654.x

Muehlroth, B. E., Sander, M. C., Fandakova, Y., Grandy, T. H., Rasch, B., Shing, Y. L., & Werkle-Bergner, M. (2019). Precise slow oscillation–spindle coupling promotes memory consolidation in younger and older adults. Scientific Reports, 9(1), Article 1940. https://doi.org/10.1038/s41598-018-36557-z

Muehlroth, B. E., Sander, M. C., Fandakova, Y., Grandy, T. H., Rasch, B., Shing, Y. L., & Werkle-Bergner, M. (2020). Memory quality modulates the effect of aging on memory consolidation during sleep: Reduced maintenance but intact gain. NeuroImage, 209, Article 116490. https://doi.org/10.1016/j.neuroimage.2019.116490

Muehlroth, B. E., & Werkle-Bergner, M. (2020). Understanding the interplay of sleep and aging: Methodological challenges. Psychophysiology, 57(3), Article e13523. https://doi.org/10.1111/psyp.13523

Muller, L., Piantoni, G., Koller, D., Cash, S. S., Halgren, E., & Sejnowski, T. J. (2016). Rotating waves during human sleep spindles organize global patterns of activity that repeat precisely through the night. eLife, 5, Article e17267. https://doi.org/10.7554/eLife.17267

Murphy, M., Riedner, B. A., Huber, R., Massimini, M., Ferrarelli, F., & Tononi, G. (2009). Source modeling sleep slow waves. Proceedings of the National Academy of Sciences of the United States of America, 106(5), 1608–1613. https://doi.org/10.1073/pnas.0807933106

Nettersheim, A., Hallschmid, M., Born, J., & Diekelmann, S. (2015). The role of sleep in motor sequence consolidation: Stabilization rather than enhancement. Journal of Neuroscience, 35(17), 6696–6702. https://doi.org/10.1523/JNEUROSCI.1236-14.2015

Niethard, N., Ngo, H.-V. V., Ehrlich, I., & Born, J. (2018). Cortical circuit activity underlying sleep slow oscillations and spindles. Proceedings of the National Academy of Sciences of the United States of America, 115(39), E9220–E9229. https://doi.org/10.1073/pnas.1805517115

Nir, Y., Staba, R. J., Andrillon, T., Vyazovskiy, V. V., Cirelli, C., Fried, I., & Tononi, G. (2011). Regional slow waves and spindles in human sleep. Neuron, 70(1), 153–169. https://doi.org/10.1016/j.neuron.2011.02.043

Nunez, P. L. (2000). Toward a quantitative description of large-scale neocortical dynamic function and EEG. Behavioral and Brain Sciences, 23(3), 371–398. https://doi.org/10.1017/S0140525X00003253

Ohayon, M., Wickwire, E. M., Hirshkowitz, M., Albert, S. M., Avidan, A., Daly, F. J., Dauvilliers, Y., Ferri, R., Fung, C., Gozal, D., Hazen, N., Krystal, A., Lichstein, K., Mallampalli, M., Plazzi, G., Rawding, R., Scheer, F. A., Somers, V., & Vitiello, M. V. (2017). National Sleep Foundation’s sleep quality recommendations: First report. Sleep Health, 3(1), 6–19. https://doi.org/10.1016/j.sleh.2016.11.006

Olbrich, E., Rusterholz, T., LeBourgeois, M. K., & Achermann, P. (2017). Developmental changes in sleep oscillations during early childhood. Neural Plasticity, 2017, Article ID 6160959. https://doi.org/10.1155/2017/6160959

Oostenveld, R., Fries, P., Maris, E., & Schoffelen, J.-M. (2011). FieldTrip: Open source software for advanced analysis of MEG, EEG, and invasive electrophysiological data. Computational Intelligence and Neuroscience, 2011, 1–9. https://doi.org/10.1155/2011/156869

Peyrache, A., Khamassi, M., Benchenane, K., Wiener, S. I., & Battaglia, F. P. (2009). Replay of rule-learning related neural patterns in the prefrontal cortex during sleep. Nature Neuroscience, 12(7), 919–926. https://doi.org/10.1038/nn.2337

Peyrache, A., & Seibt, J. (2020). A mechanism for learning with sleep spindles. Philosophical Transactions of the Royal Society B: Biological Sciences, 375(1799), Article 20190230. https://doi.org/10.1098/rstb.2019.0230

Piantoni, G., Astill, R. G., Raymann, R. J. E. M., Vis, J. C., Coppens, J. E., & Van Someren, E. J. W. (2013). Modulation of gamma and spindle-range power by slow oscillations in scalp sleep EEG of children. International Journal of Psychophysiology, 89(2), 252–258. https://doi.org/10.1016/j.ijpsycho.2013.01.017

Piantoni, G., Poil, S.-S., Linkenkaer-Hansen, K., Verweij, I. M., Ramautar, J. R., Van Someren, E. J. W., & Van Der Werf, Y. D. (2013). Individual differences in white matter diffusion affect sleep oscillations. Journal of Neuroscience, 33(1), 227–233. https://doi.org/10.1523/JNEUROSCI.2030-12.2013

Prehn-Kristensen, A., Göder, R., Fischer, J., Wilhelm, I., Seeck-Hirschner, M., Aldenhoff, J., & Baving, L. (2011). Reduced sleep-associated consolidation of declarative memory in attention-deficit/hyperactivity disorder. Sleep Medicine, 12(7), 672–679. https://doi.org/10.1016/j.sleep.2010.10.010

Prehn-Kristensen, A., Munz, M., Göder, R., Wilhelm, I., Korr, K., Vahl, W., Wiesner, C. D., & Baving, L. (2014). Transcranial oscillatory direct current stimulation during sleep improves declarative memory consolidation in children with attention-deficit/hyperactivity disorder to a level comparable to healthy controls. Brain Stimulation, 7(6), 793–799. https://doi.org/10.1016/j.brs.2014.07.036

Purcell, S. M., Manoach, D. S., Demanuele, C., Cade, B. E., Mariani, S., Cox, R., Panagiotaropoulou, G., Saxena, R., Pan, J. Q., Smoller, J. W., Redline, S., & Stickgold, R. (2017). Characterizing sleep spindles in 11,630 individuals from the National Sleep Research Resource. Nature Communications, 8(1), 15930. https://doi.org/10.1038/ncomms15930

Rasch, B., & Born, J. (2013). About sleep’s role in memory. Physiological Reviews, 93(2), 681–766. https://doi.org/10.1152/physrev.00032.2012

Rosanova, M., & Ulrich, D. (2005). Pattern-specific associative long-term potentiation induced by a sleep spindle-related spike train. Journal of Neuroscience, 25(41), 9398–9405. https://doi.org/10.1523/JNEUROSCI.2149-05.2005

Saletin, J. M., van der Helm, E., & Walker, M. P. (2013). Structural brain correlates of human sleep oscillations. NeuroImage, 83, 658–668. https://doi.org/10.1016/j.neuroimage.2013.06.021

Schabus, M., Dang-Vu, T. T., Albouy, G., Balteau, E., Boly, M., Carrier, J., Darsaud, A., Degueldre, C., Desseilles, M., Gais, S., Phillips, C., Rauchs, G., Schnakers, C., Sterpenich, V., Vandewalle, G., Luxen, A., & Maquet, P. (2007). Hemodynamic cerebral correlates of sleep spindles during human non-rapid eye movement sleep. Proceedings of the National Academy of Sciences of the United States of America, 104(32), 13164–13169. https://doi.org/10.1073/pnas.0703084104

Schabus, M., Gruber, G., Parapatics, S., Sauter, C., Klösch, G., Anderer, P., Klimesch, W., Saletu, B., & Zeitlhofer, J. (2004). Sleep spindles and their significance for declarative memory consolidation. Sleep, 27(8), 1479–1485. https://doi.org/10.1093/sleep/27.7.1479

Schapiro, A. C., McDevitt, E. A., Rogers, T. T., Mednick, S. C., & Norman, K. A. (2018). Human hippocampal replay during rest prioritizes weakly learned information and predicts memory performance. Nature Communications, 9(1), Article 3920. https://doi.org/10.1038/s41467-018-06213-1

Schlarb, A. A., Schwerdtle, B., & Hautzinger, M. (2010). Validation and psychometric properties of the German version of the Children’s Sleep Habits Questionnaire (CSHQ-DE). Somnologie – Schlafforschung und Schlafmedizin, 14(4), 260–266. https://doi.org/10.1007/s11818-010-0495-4

Schneider, W., & Sodian, B. (1997). Memory strategy development: Lessons from longitudinal research. Developmental Review, 17(4), 442–461. https://doi.org/10.1006/drev.1997.0441

Schoch, S. F., Cordi, M. J., & Rasch, B. (2017). Modulating influences of memory strength and sensitivity of the retrieval test on the detectability of the sleep consolidation effect. Neurobiology of Learning and Memory, 145, 181–189. https://doi.org/10.1016/j.nlm.2017.10.009

Scholle, S., Zwacka, G., & Scholle, H. C. (2007). Sleep spindle evolution from infancy to adolescence. Clinical Neurophysiology, 118(7), 1525–1531. https://doi.org/10.1016/j.clinph.2007.03.007

Schreiner, T., & Rasch, B. (2018). To gain or not to gain: The complex role of sleep for memory. Cortex, 101, 282–287. https://doi.org/10.1016/j.cortex.2016.06.011

Schwerdtle, B., Kanis, Kahl, Kübler, & Schlarb, A. (2012). Children’s Sleep Comic: Development of a new diagnostic tool for children with sleep disorders. Nature and Science of Sleep, 2012(4), 97– 102. https://doi.org/10.2147/NSS.S33127

Shing, Y. L., Werkle-Bergner, M., Li, S.-C., & Lindenberger, U. (2008). Associative and strategic components of episodic memory: A life-span dissociation. Journal of Experimental Psychology: General, 137(3), 495–513. https://doi.org/10.1037/0096-3445.137.3.495

Shinomiya, S., Nagata, K., Takahashi, K., & Masumura, T. (1999). Development of sleep spindles in young children and adolescents. Clinical Electroencephalography, 30(2), 39–43. https://doi.org/10.1177/155005949903000203

Siapas, A. G., & Wilson, M. A. (1998). Coordinated interactions between hippocampal ripples and cortical spindles during slow-wave sleep. Neuron, 21(5), 1123–1128. https://doi.org/10.1016/S0896-6273(00)80629-7

Sirota, A., Csicsvari, J., Buhl, D., & Buzsaki, G. (2003). Communication between neocortex and hippocampus during sleep in rodents. Proceedings of the National Academy of Sciences of the United States of America, 100(4), 2065–2069. https://doi.org/10.1073/pnas.0437938100

Staresina, B. P., Bergmann, T. O., Bonnefond, M., van der Meij, R., Jensen, O., Deuker, L., Elger, C. E., Axmacher, N., & Fell, J. (2015). Hierarchical nesting of slow oscillations, spindles and ripples in the human hippocampus during sleep. Nature Neuroscience, 18(11), 1679–1686. https://doi.org/10.1038/nn.4119

Steriade, M. (2006). Grouping of brain rhythms in corticothalamic systems. Neuroscience, 137(4), 1087–1106. https://doi.org/10.1016/j.neuroscience.2005.10.029

Steriade, M., Contreras, D., Curro Dossi, R., & Nunez, A. (1993). The slow (< 1 Hz) oscillation in reticular thalamic and thalamocortical neurons: Scenario of sleep rhythm generation in interacting thalamic and neocortical networks. Journal of Neuroscience, 13(8), 3284–3299. https://doi.org/10.1523/JNEUROSCI.13-08-03284.1993

Steriade, M., & Llinás, R. R. (1988). The functional states of the thalamus and the associated neuronal interplay. Physiological Reviews, 68(3), 649–742. https://doi.org/10.1152/physrev.1988.68.3.649

Steriade, M., Nunez, A., & Amzica, F. (1993). A novel slow (<1 Hz) oscillation of neocortical neurons in vivo: Depolarizing and hyperpolarizing components. Journal of Neuroscience, 13(8), 3252– 3265. https://doi.org/10.1523/JNEUROSCI.13-08-03252.1993.

Tarokh, L., & Carskadon, M. A. (2010). Developmental changes in the human sleep EEG during early adolescence. Sleep, 33(6), 801–809. https://doi.org/10.1093/sleep/33.6.801

Timofeev, I., & Chauvette, S. (2013). The spindles: Are they still thalamic? Sleep, 36(6), 825–826. https://doi.org/10.5665/sleep.2702

Timofeev, I., Grenier, F., Bazhenov, M., Houweling, A. R., Sejnowski, T. J., & Steriade, M. (2002). Short- and medium-term plasticity associated with augmenting responses in cortical slabs and spindles in intact cortex of cats *in vivo*. Journal of Physiology, 542(2), 583–598. https://doi.org/10.1113/jphysiol.2001.013479

Timofeev, I., Schoch, S. F., LeBourgeois, M. K., Huber, R., Riedner, B. A., & Kurth, S. (2020). Spatio-temporal properties of sleep slow waves and implications for development. Current Opinion in Physiology, 15, 172–182. https://doi.org/10.1016/j.cophys.2020.01.007

Tucker, M. A., & Fishbein, W. (2008). Enhancement of declarative memory performance following a daytime nap is contingent on strength of initial task acquisition. Sleep, 31(2), 197–203. https://doi.org/10.1093/sleep/31.2.197

Tulving, E. (1967). The effects of presentation and recall of material in free-recall learning. Journal of Verbal Learning and Verbal Behavior, 6(2), 175–184. https://doi.org/10.1016/S0022-5371(67)80092-6

Tulving, E., & Pearlstone, Z. (1966). Availability versus accessibility of information in memory for words. Journal of Verbal Learning and Verbal Behavior, 5(4), 381–391. https://doi.org/10.1016/S0022-5371(66)80048-8

Tulving, E., & Psotka, J. (1971). Retroactive inhibition in free recall: Inaccessibility of information available in the memory store. Journal of Experimental Psychology, 87(1), 1–8. https://doi.org/10.1037/h0030185

Ujma, P. P., Gombos, F., Genzel, L., Konrad, B. N., Simor, P., Steiger, A., Dresler, M., & Bódizs, R. (2015). A comparison of two sleep spindle detection methods based on all night averages: Individually adjusted vs. fixed frequencies. Frontiers in Human Neuroscience, 9, Article 52. https://doi.org/10.3389/fnhum.2015.00052

Urbain, C., De Tiège, X., Op De Beeck, M., Bourguignon, M., Wens, V., Verheulpen, D., Van Bogaert, P., & Peigneux, P. (2016). Sleep in children triggers rapid reorganization of memory-related brain processes. NeuroImage, 134, 213–222. https://doi.org/10.1016/j.neuroimage.2016.03.055

Wang, J.-Y., Weber, F. D., Zinke, K., Noack, H., & Born, J. (2017). Effects of sleep on word pair memory in children: Separating item and source memory aspects. Frontiers in Psychology, 8, Article 1533. https://doi.org/10.3389/fpsyg.2017.01533

Watson, B. O., & Buzsáki, G. (2015). Sleep, memory & brain rhythms. Daedalus, 144(1), 67–82. https://doi.org/10.1162/DAED_a_00318

Werner, H., LeBourgeois, M. K., Geiger, A., & Jenni, O. G. (2009). Assessment of chronotype in four- to elevenyYear-old children: Reliability and validity of the Children’s ChronoType Questionnaire (CCTQ). Chronobiology International, 26(5), 992–1014. https://doi.org/10.1080/07420520903044505

Werth, E., Achermann, P., Dijk, D.-J., & Borbely, A. A. (1997). Spindle frequency activity in the sleep EEG: individual differences and topographic distribution. Electroencephalography and Clinical Neurophysiology, 103(5), 535–554. https://doi.org/10.1016/s0013-4694(97)00070-9

Wilhelm, I., Diekelmann, S., & Born, J. (2008). Sleep in children improves memory performance on declarative but not procedural tasks. Learning & Memory, 15(5), 373–377. https://doi.org/10.1101/lm.803708

Wilhelm, I., Metzkow-Mészàros, M., Knapp, S., & Born, J. (2012). Sleep-dependent consolidation of procedural motor memories in children and adults: The pre-sleep level of performance matters: Sleep-dependent consolidation of motor memories. Developmental Science, 15(4), 506–515. https://doi.org/10.1111/j.1467-7687.2012.01146.x

Wilhelm, I., Rose, M., Imhof, K. I., Rasch, B., Büchel, C., & Born, J. (2013). The sleeping child outplays the adult’s capacity to convert implicit into explicit knowledge. Nature Neuroscience, 16(4), 391–393. https://doi.org/10.1038/nn.3343

Wilhelm, I., Schreiner, T., Beck, J., & Rasch, B. (2020). No effect of targeted memory reactivation during sleep on retention of vocabulary in adolescents. Scientific Reports, 10(1), Article 4255. https://doi.org/10.1038/s41598-020-61183-z

Wilson, M., & McNaughton, B. (1994). Reactivation of hippocampal ensemble memories during sleep. Science, 265(5172), 676–679. https://doi.org/10.1126/science.8036517

