## Supplemental Information for "Electrophysiological Indicators of Sleep-associated Memory Consolidation in 5- to 6-Year-Old Children"

Table S1.

*Results from the Wilcoxon signed-rank tests comparing indicators of sleep quality between the two PSG nights.*

| | Baseline Night | | | Learning Night | | | | | |
|---|---|---|---|---|---|---|---|---|---|
| Variables | *Median* | *1.Quartile* | *3.Quartile* | *Median* | *1.Quartile* | *3.Quartile* | *Z* | *p* | $CI_{2.5, 97.5}$ |
| TST (min) | 586.50 | 549.50 | 599.00 | 591.00 | 552.62 | 605.75 | -0.65 | 0.516 | [-2.54, -0.02] |
| N1 (%) | 8.70 | 7.17 | 11.61 | 8.41 | 5.71 | 10.63 | -0.60 | 0.546 | [-2.65, -0.01] |
| N2 (%) | 48.42 | 42.76 | 51.11 | 47.85 | 44.46 | 50.78 | -0.92 | 0.360 | [-2.67, -0.04] |
| N3 (%) | 18.16 | 14.06 | 21.99 | 18.42 | 14.92 | 20.82 | -1.03 | 0.303 | [-2.84, -0.07] |
| NREM (%) | 66.14 | 62.96 | 69.90 | 65.95 | 63.33 | 68.41 | -0.01 | 0.989 | [-2.22, -0.01] |
| REM (%) | 23.82 | 20.58 | 28.58 | 25.44 | 22.87 | 27.62 | -0.89 | 0.375 | [-2.73, -0.04] |
| WASO (min) | 1.90 | 0.66 | 11.81 | 0.58 | 0.31 | 2.49 | -1.90 | 0.057 | [-3.45, -0.23] |

*Note*. *Z*- and *p*-values were derived from non-parametric Wilcoxon signed-rank tests comparing indicators of sleep quality between the baseline and the learning night. The confidence interval ($CI_{2.5, 97.5}$) represents the 95-% simple bootstrap percentile confidence interval of the *Z*-value, based on 5000 case re-samples. TST: total sleep time; NREM: non-rapid eye movement sleep; REM: rapid eye movement sleep; WASO: wake after sleep onset.

Table S2.

***(A)*** *Overview of results from the mixed factorial ANOVA examining recall performance between the different RECALL SESSIONs (Recall & Feedback, Evening Recall, Morning Recall) and GROUPs (Group$_{50}$, Group$_{100}$).* ***(B)*** *Results from the post-hoc Wilcoxon signed-rank tests for the main effect of RECALL SESSION.*

| A | | | | | | |
|---|---|---|---|---|---|---|
| Predictor | $df_{Num}$ | $df_{Den}$ | *Epsilon* | *F* | *p* | $\eta^2_G$ |
| Group | 1.00 | 22.00 | | 1.08 | 0.309 | 0.04 |
| Recall Session | 1.16 | 25.56 | 0.58 | 44.47 | <0.001 | 0.15 |
| Interaction | 1.16 | 25.56 | 0.58 | 0.53 | 0.501 | 0.00 |

*Note.* $df_{Num}$ indicates degrees of freedom numerator. $df_{Den}$ indicates degrees of freedom denominator. Epsilon indicates Greenhouse-Geisser multiplier for degrees of freedom, *p*-values and degrees of freedom in the table incorporate this correction. $\eta^2_G$ indicates generalised eta-squared.

| B | | | |
|---|---|---|---|
| Recall Session Contrast | *Z* | *p* | $CI_{2.5, 97.5}$ |
| Recall & Feedback vs. Evening Recall | -3.96 | <0.001 | [-4.13, -2.83] |
| Recall & Feedback vs. Morning Recall | -3.98 | <0.001 | [-4.13, -2.89] |
| Evening Recall vs. Morning Recall | -0.14 | 0.887 | [-2.23, 0.02] |

*Note.* *Z*- and *p*-values were derived from non-parametric Wilcoxon signed-rank tests comparing percentage recall performance between recall sessions. The confidence interval ($CI_{2.5, 97.5}$) represents the 95-% simple bootstrap percentile confidence interval of the *Z*-value, based on 5000 case re-samples.

Table S3.

*Overview of descriptive measures of “high” and individually identified SPs for averaged frontal and centro-parietal recording sites during the learning night.*

| | | SP Type | | | | | |
|---|---|---|---|---|---|---|---|
| | | “High” SPs | | | Individually Identified SPs | | |
| Measure | Topography | *Median* | *1.Quartile* | *3.Quartile* | *Median* | *1.Quartile* | *3.Quartile* |
| Frequency | Frontal | 13.13 | 12.88 | 13.54 | 11.08 | 10.73 | 11.40 |
| | Centro-parietal | 13.56 | 13.25 | 13.91 | 11.84 | 11.52 | 12.09 |
| Density | Frontal | 0.54 | 0.36 | 0.68 | 1.38 | 0.97 | 1.62 |
| | Centro-parietal | 0.53 | 0.45 | 0.69 | 0.91 | 0.78 | 1.17 |
| Amplitude | Frontal | 17.35 | 14.00 | 19.20 | 34.98 | 29.50 | 47.44 |
| | Centro-parietal | 12.55 | 11.70 | 14.34 | 17.41 | 15.68 | 20.65 |

Table S4.

***(A)** Results from the post-hoc Wilcoxon signed-rank tests on the main effect of "ELECTRODE" contrasting individually identified frontal sleep spindle frequency, density, and amplitude with measures derived from central and parietal electrodes. **(B)** Results from the Wilcoxon signed-rank tests comparing the averaged individually identified frontal and centro-parietal sleep spindle frequency, density, and amplitude. Results show that frontal and centro-parietal SPs differed between averaged frontal and centro-parietal sites.*

| A | | | | |
|---|---|---|---|---|
| SP measure | Electrode Contrast | *Z* | *p* | $CI_{2.5, 97.5}$ |
| Frequency | F3 vs. C3 | -5.30 | <0.001 | [-4.10, -4.09] |
| | F3 vs. Cz | -5.30 | <0.001 | [-4.10, -4.09] |
| | F3 vs.C4 | -5.17 | <0.001 | [-4.10, -3.90] |
| | F3 vs. Pz | -5.09 | <0.001 | [-4.10, -3.74] |
| | F4 vs.C3 | -5.30 | <0.001 | [-4.10, -4.09] |
| | F4 vs. Cz | -5.30 | <0.001 | [-4.10, -4.09] |
| | F4 vs.C4 | -5.17 | <0.001 | [-4.10, -3.90] |
| | F4 vs. Pz | -5.09 | <0.001 | [-4.10, -3.74] |
| Density | F3 vs.C3 | -3.75 | <0.001 | [-3.91, -2.00] |
| | F3 vs. Cz | -2.10 | 0.040 | [-3.48, -0.17] |
| | F3 vs.C4 | -4.90 | <0.001 | [-4.09, -3.49] |
| | F3 vs. Pz | -3.71 | <0.001 | [-3.76, -1.89] |
| | F4 vs.C3 | -3.49 | <0.001 | [-3.80, -1.50] |
| | F4 vs. Cz | -2.27 | 0.020 | [-3.51, -0.38] |
| | F4 vs.C4 | -4.86 | <0.001 | [-4.10, -3.48] |
| | F4 vs. Pz | -3.24 | 0.001 | [-3.70, -1.35] |
| Amplitude | F3 vs.C3 | -5.04 | <0.001 | [-4.10, -3.60] |
| | F3 vs. Cz | -5.04 | <0.001 | [-4.10, -3.60] |
| | F3 vs.C4 | -5.04 | <0.001 | [-4.10, -3.60] |
| | F3 vs. Pz | -4.76 | <0.001 | [-4.00, -3.27] |
| | F4 vs.C3 | -5.17 | <0.001 | [-4.10, -3.81] |
| | F4 vs. Cz | -5.04 | <0.001 | [-4.10, -3.68] |
| | F4 vs.C4 | -5.17 | <0.001 | [-4.10, -3.81] |
| | F4 vs. Pz | -4.90 | <0.001 | [-4.10, -3.49] |

*Note. Z*- and *p*-values were derived from non-parametric Wilcoxon signed-rank tests comparing the respective SP measure between recording sites (electrode contrast). The confidence interval ($CI_{2.5, 97.5}$) represents the 95-% simple bootstrap percentile confidence interval of the *Z*-value, based on 5000 case re-samples.

| B | | | | |
|---|---|---|---|---|
| SP Measure | Topography Contrast | *Z* | *p* | $CI_{2.5, 97.5}$ |
| Frequency | Frontal vs. Centro-parietal | -5.30 | <0.001 | [-4.28, -4.27] |
| Density | Frontal vs. Centro-parietal | -4.17 | <0.001 | [-4.00, -2.50] |
| Amplitude | Frontal vs. Centro-parietal | -5.30 | <0.001 | [-4.10, -4.09] |

*Note. Z*- and *p*-values were derived from non-parametric Wilcoxon signed-rank tests comparing the respective SP measure between averaged frontal and centro-parietal sites. The confidence interval ($CI_{2.5, 97.5}$) represents the 95-% simple bootstrap percentile confidence interval of the *Z*-value, based on 5000 case re-samples.

Table S5.

*Overview of results from the post-hoc Wilcoxon signed-rank tests examining differences between the co-occurrence of SP centres of the two individually identified SP types (slow frontal vs. fast centro-parietal) with SOs from different topographical locations (frontal, centro-parietal, occipital) during the learning night.* ***(A)*** *Contrasts comparing the occurrence of individually identified slow frontal and fast centro-parietal SPs between SOs in different locations.* ***(B)*** *Contrasts comparing occurrence of individually identified slow frontal against individually identified fast centro-parietal SPs during frontal, centro-parietal, and occipital SOs during the learning night.*

| A | | | | |
|---|---|---|---|---|
| SP Type | SO Topography Contrast | *Z* | *p* | $CI_{2.5, 97.5}$ |
| Slow Frontal | Frontal vs. Centro-parietal | -3.48 | <0.001 | [-3.93, -1.63] |
| Slow Frontal | Frontal vs. Occipital | -5.17 | <0.001 | [-4.10, -3.90] |
| Slow Frontal | Centro-parietal vs. Occipital | -5.30 | <0.001 | [-4.10, -4.09] |
| Fast Centro-parietal | Frontal vs. Centro-parietal | -4.22 | <0.001 | [-4.10, -2.60] |
| Fast Centro-parietal | Frontal vs. Occipital | -5.10 | <0.001 | [-4.10, -3.71] |
| Fast Centro-parietal | Centro-parietal vs. Occipital | -5.30 | <0.001 | [-4.10, -4.09] |

*Note. Z*- and *p*-values were derived from non-parametric Wilcoxon signed-rank tests comparing individually identified slow frontal and fast centro-parietal SP occurrence during SOs in different topographical locations (SO topography contrast).The confidence interval ($CI_{2.5, 97.5}$) represents the 95-% simple bootstrap percentile confidence interval of the *Z*-value, based on 5000 case re-samples.

| B | | | | |
|---|---|---|---|---|
| SO Topography | SP Type Contrast | *Z* | *p* | $CI_{2.5, 97.5}$ |
| Frontal | Slow Frontal vs. Fast Centro-parietal | -4.36 | <0.001 | [-4.10, -2.73] |
| Centro-parietal | Slow Frontal vs. Fast Centro-parietal | -3.52 | <0.001 | [-3.96, -1.56] |
| Occipital | Slow Frontal vs. Fast Centro-parietal | -5.30 | <0.001 | [-4.10, -3.25] |

*Note. Z*- and *p*-values were derived from non-parametric Wilcoxon signed-rank tests comparing individually identified slow frontal with individually identified fast centro-parietal SPs (SP type contrast) during SOs in different topographical locations. The confidence interval ($CI_{2.5, 97.5}$) represents the 95-% simple bootstrap percentile confidence interval of the *Z*-value, based on 5000 case re-samples.

Table S6.

***(A)*** *Overview of results from the repeated measure ANOVA examining the co-occurrence of SO DOWN peaks in different topographical locations with individually identified slow frontal and fast centro-parietal SPs during the learning night with the within-person factors SP TYPE (slow frontal, fast centro-parietal) and SO TOPOGRAPHY (frontal, centro-parietal, occipital).* ***(B)*** *Results from the post-hoc Wilcoxon signed-rank tests on the main effects "SO Topography" and "SP Type".*

| A | | | | | | |
|---|---|---|---|---|---|---|
| Predictor | $df_{Num}$ | $df_{Den}$ | *Epsilon* | *F* | *p* | $\eta^2_G$ |
| SP Type | 1.00 | 23.00 | | 36.55 | <0.001 | 0.25 |
| SO Topography | 1.48 | 34.07 | 0.74 | 7.51 | 0.004 | 0.03 |
| Interaction | 1.40 | 32.31 | 0.70 | 1.23 | 0.293 | 0.00 |

*Note.* $df_{Num}$ indicates degrees of freedom numerator. $df_{Den}$ indicates degrees of freedom denominator. Epsilon indicates Greenhouse-Geisser multiplier for degrees of freedom, *p*-values and degrees of freedom in the table incorporate this correction. $\eta^2_G$ indicates generalised eta-squared.

| B | | | | |
|---|---|---|---|---|
| Main Effect | Contrast | *Z* | *p* | $CI_{2.5, 97.5}$ |
| SO Topography | Frontal vs. Centro-parietal | -3.71 | <0.001 | [-4.10, -2.31] |
| | Frontal vs. Occipital | -0.21 | 0.833 | [-2.34, -0.03] |
| | Centro-parietal vs. Occipital | -2.24 | 0.025 | [-4.10, -1.40] |
| SP Type | Slow Frontal vs. Fast Centro-parietal | -4.86 | <0.001 | [-4.86, -3.41] |

*Note. Z-* and *p*-values were derived from non-parametric Wilcoxon signed-rank for the main effects of the repeated measure ANOVA comparing SO DOW -peak occurrence in different topographical locations during individually identified slow frontal and fast centro-parietal SPs. The confidence interval ($CI_{2.5, 97.5}$) represents the 95-% simple bootstrap percentile confidence interval of the *Z*-value, based on 5000 case re-samples.

*Table S7.*

***(A)*** *Overview of results from the repeated measure ANOVA examining the co-occurrence of "high"SP centres with SOs during the learning night in different topographical locations with the within-person factors SP TYPE ("high" frontal, "high" centro-parietal) and SO TOPOGRAPHY (frontal, centro-parietal, occipital).* ***(B)*** *Results from the post-hoc Wilcoxon signed-rank tests on the interaction effect.*

| A | | | | | | |
|---|---|---|---|---|---|---|
| Predictor | $df_{Num}$ | $df_{Den}$ | *Epsilon* | *F* | *p* | $\eta^2_G$ |
| SP Type | 1.00 | 23.00 | | 1.82 | 0.190 | 0.01 |
| SO Topography | 1.50 | 34.57 | 0.75 | 92.27 | <0.001 | 0.46 |
| Interaction | 1.82 | 41.94 | 0.91 | 6.83 | 0.003 | 0.01 |

*Note.* $df_{Num}$ indicates degrees of freedom numerator. $df_{Den}$ indicates degrees of freedom denominator. Epsilon indicates Greenhouse-Geisser multiplier for degrees of freedom, *p*-values and degrees of freedom in the table incorporate this correction. $\eta^2_G$ indicates generalised eta-squared.

| B | | | | |
|---|---|---|---|---|
| SP Type | SO Topography Contrast | *Z* | *p* | $CI_{2.5, 97.5}$ |
| "High" Frontal | Frontal vs. Centro-parietal | -3.45 | <0.001 | [-3.97, -1.50] |
| "High" Frontal | Frontal vs. Occipital | -4.99 | <0.001 | [-4.10, -3.61] |
| "High" Frontal | Centro-parietal vs. Occipital | -5.17 | <0.001 | [-4.10, -3.90] |
| "High" Centro-parietal | Frontal vs. Centro-parietal | -4.41 | <0.001 | [-4.09, -2.79] |
| "High" Centro-parietal | Frontal vs. Occipital | -4.99 | <0.001 | [-4.10, -3.61] |
| "High" Centro-parietal | Centro-parietal vs. Occipital | -5.30 | <0.001 | [-4.10, -4.09] |

*Note. Z*- and *p*-values were derived from non-parametric Wilcoxon signed-rank tests comparing "high" frontal or "high" centro-parietal SP occurrence during SOs in different topographical locations. The confidence interval ($CI_{2.5, 97.5}$) represents the 95-% simple bootstrap percentile confidence interval of the *Z*-value, based on 5000 case re-samples.

Table S8.

***(A)** Overview of results from the repeated measure ANOVA examining the co-occurrence of SO DOWN peaks in different topographical locations with "high" SPs during the learning night with the within-person factors SP TYPE ("high" frontal, "high" centro-parietal) and SO TOPOGRAPHY (frontal, centro-parietal, occipital). **(B)** Results from the post-hoc Wilcoxon signed-rank tests on interaction effect.*

| A | | | | | | |
|---|---|---|---|---|---|---|
| Predictor | $df_{Num}$ | $df_{Den}$ | *Epsilon* | *F* | *p* | $\eta^2_G$ |
| SP Type | 1.00 | 23.00 | | 0.07 | 0.791 | 0.00 |
| SO Topography | 1.91 | 43.87 | 0.95 | 24.12 | <0.001 | 0.08 |
| Interaction | 1.57 | 36.15 | 0.79 | 6.82 | 0.006 | 0.01 |

*Note.* $df_{Num}$ indicates degrees of freedom numerator. $df_{Den}$ indicates degrees of freedom denominator. Epsilon indicates Greenhouse-Geisser multiplier for degrees of freedom, *p*-values and degrees of freedom in the table incorporate this correction. $\eta^2_G$ indicates generalised eta-squared.

| B | | | | |
|---|---|---|---|---|
| SP Type | SO Topography Contrast | *Z* | *p* | $CI_{2.5, 97.5}$ |
| "High" Frontal | Frontal vs. Centro-parietal | -3.48 | <0.001 | [-4.06, -1.72] |
| "High" Frontal | Frontal vs. Occipital | -1.26 | 0.208 | [-2.89, -0.03] |
| "High" Frontal | Centro-parietal vs. Occipital | -2.53 | 0.011 | [-3.87, -0.91] |
| "High" Centro-parietal | Frontal vs. Centro-parietal | -5.09 | <0.001 | [-4.10, -3.74] |
| "High" Centro-parietal | Frontal vs. Occipital | -0.95 | 0.345 | [-2.89, -0.03] |
| "High" Centro-parietal | Centro-parietal vs. Occipital | -5.17 | <0.001 | [-4.10, -3.90] |

*Note. Z*- and *p*-values were derived from non-parametric Wilcoxon signed-rank tests comparing SO occurrence in different topographical locations during "high" frontal and centro-parietal SPs. The confidence interval ($CI_{2.5, 97.5}$) represents the 95-% simple bootstrap percentile confidence interval of the *Z*-value, based on 5000 case re-samples.

Table S9.

*Overview of results from the robust regression on memory consolidation of medium-quality memories.*

| Predictor | *β* | $CI_{2.5, 97.5}$ | *F* | *p* |
|---|---|---|---|---|
| Intercept | 0.004 | [-0.36, 0.38] | <0.001 | 0.976 |
| Age | 0.36 | [-0.02, 0.81] | 6.45 | 0.020 |
| Slow Frontal SP Density Change | 0.55 | [0.15, 0.85] | 16.44 | < 0.001 |
| Fast Centro-parietal SP Density Change | 0.09 | [-0.17, 0.65] | 0.48 | 0.495 |
| SO Amplitude | 0.29 | [0.04, 0.54] | 4.98 | 0.038 |

*Note.* The dependent variable was medium-quality memory consolidation. The confidence interval ($CI_{2.5, 97.5}$) represents the 95-% simple bootstrap percentile confidence interval of the *β*-value, based on 5000 case re-samples.

Table S10.

*Overview of results from the robust regression on memory consolidation of high-quality memories.*

| Predictor | *β* | $CI_{2.5, 97.5}$ | *F* | *p* |
|---|---|---|---|---|
| Intercept | 0.22 | [-0.21, 0.38] | 9.57 | 0.006 |
| Age | -0.02 | [-0.18, 0.35] | 0.05 | 0.822 |
| Slow Frontal SP Density Change | 0.07 | [-0.24, 0.23] | 0.79 | 0.386 |
| Fast Centro-parietal SP Density Change | 0.04 | [-0.11, 0. 57] | 0.22 | 0.641 |
| SO Amplitude | 0.12 | [0, 0.44] | 2.69 | 0.117 |

*Note.* The dependent variable was high-quality memory consolidation. The confidence interval ($CI_{2.5, 97.5}$) represents the 95-% simple bootstrap percentile confidence interval of the *β*-value, based on 5000 case re-samples.

Table S11.

*Overview of results from the robust regression on memory consolidation of low-quality memories.*

| Predictor | *β* | $CI_{2.5, 97.5}$ | *F* | *p* |
|---|---|---|---|---|
| Intercept | -0.09 | [-0.44, 0.35] | 0.33 | 0.571 |
| Age | 0.17 | [-0.15, 0.58] | 1.11 | 0.305 |
| Slow Frontal SP Density Change | -0.26 | [-0.60, 0.22] | 2.70 | 0.117 |
| Fast Centro-parietal SP Density Change | 0.39 | [-0.03, 1.10] | 5.28 | 0.033 |
| SO Amplitude | -0.23 | [-0.81, 0.10] | 2.23 | 0.152 |

*Note.* The dependent variable was low-quality memory consolidation. The confidence interval ($CI_{2.5, 97.5}$) represents the 95-% simple bootstrap percentile confidence interval of the *β*-value, based on 5000 case re-samples.

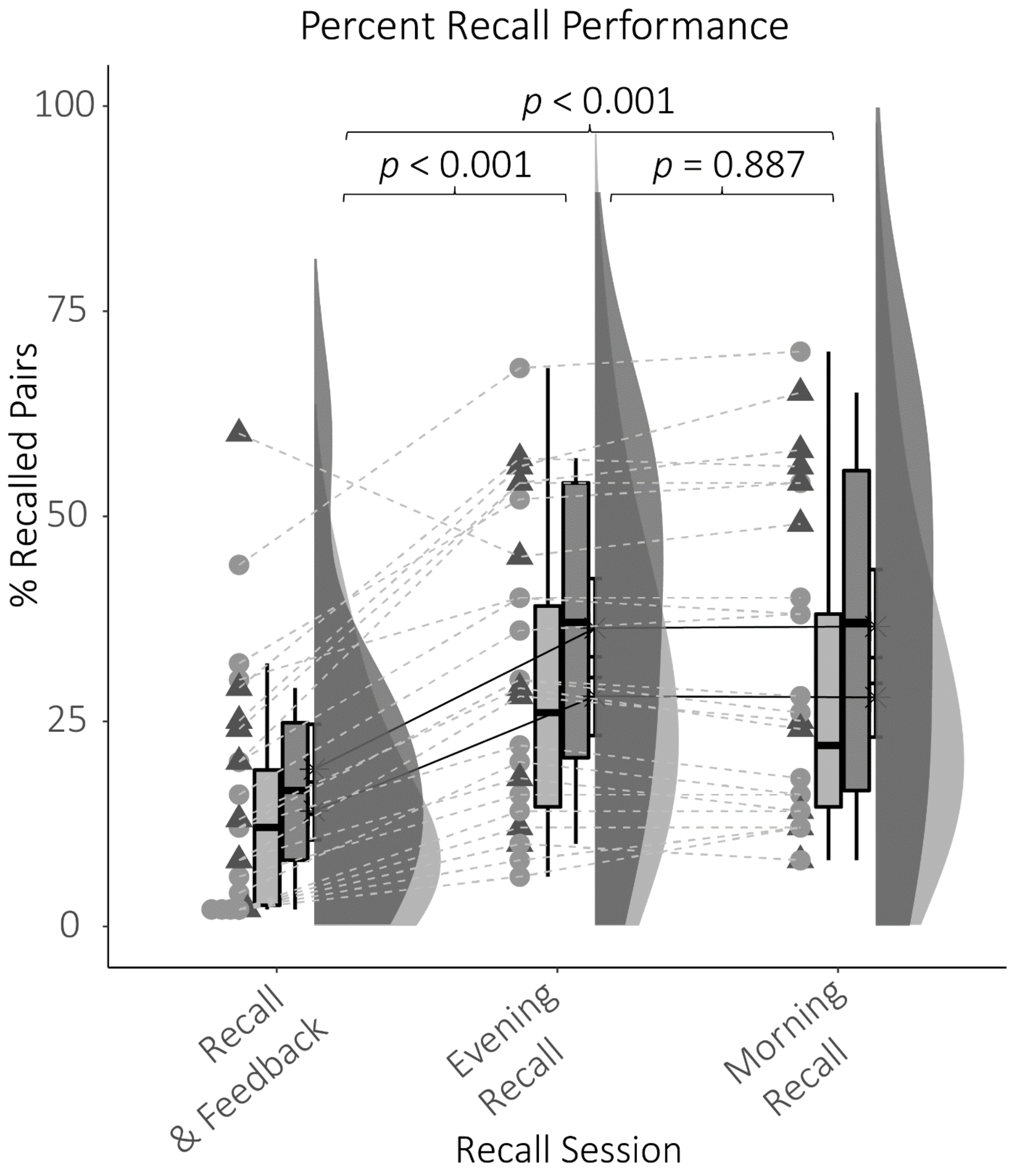


*Figure S1.* Recall performance during the three recall sessions. There was no difference between $Group_{50}$ and $Group_{100}$. Performance for both groups increased significantly from the feedback & recall session to the evening recall. Children recalled on average 31.46 % scene-object pairs during the evening recall. The percentage of recalled scene-object pairs did not differ between the evening and morning recall.

### Raw EEG Traces

**A**

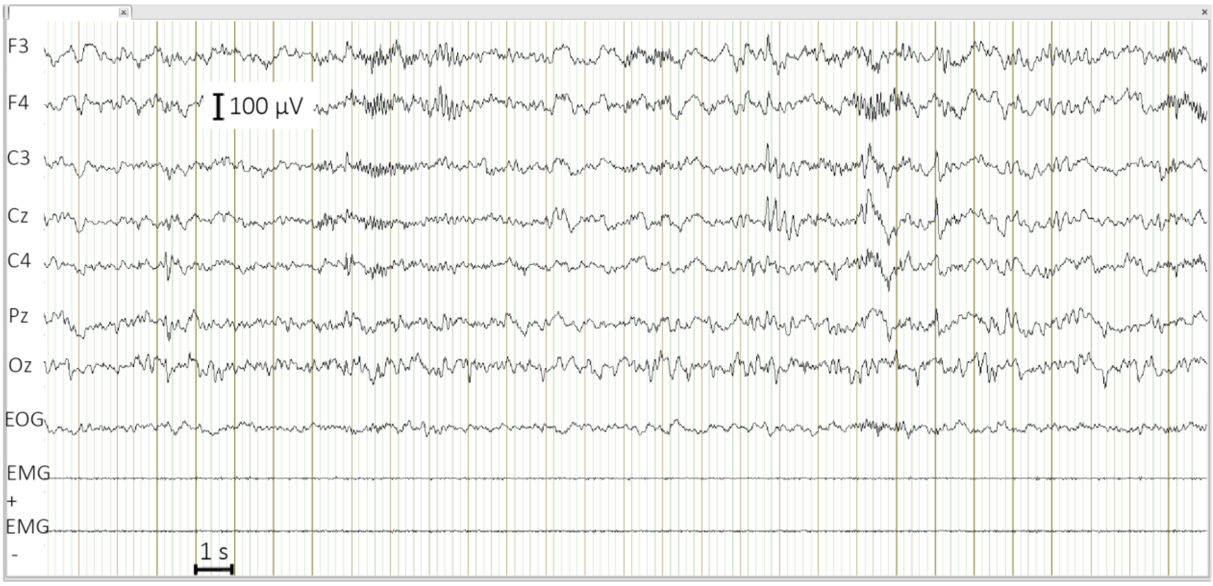


**B**

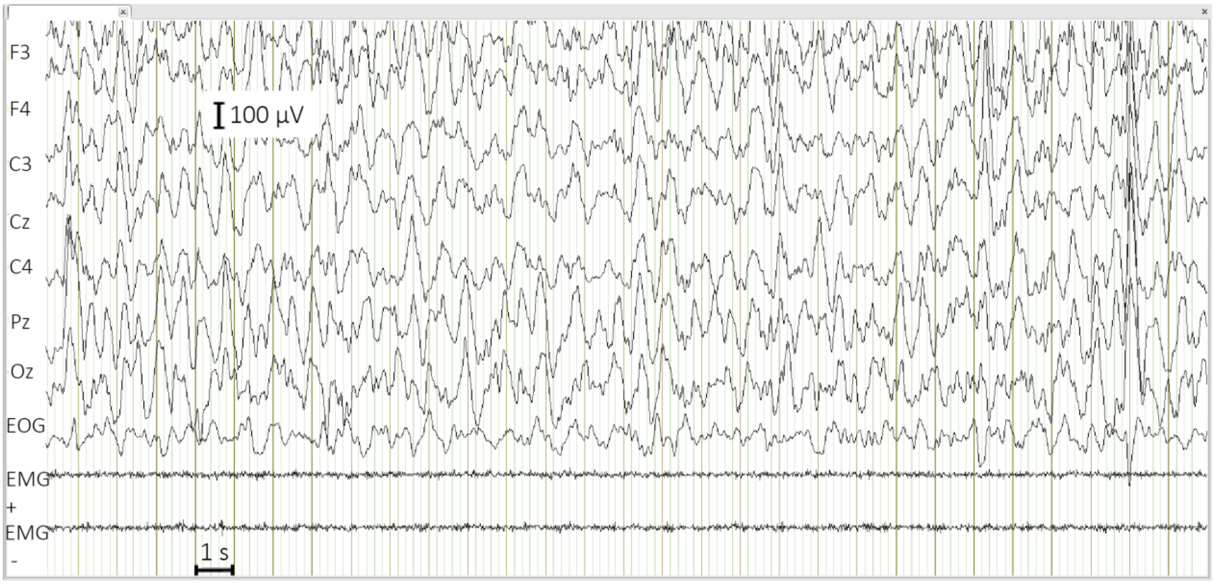

**C**

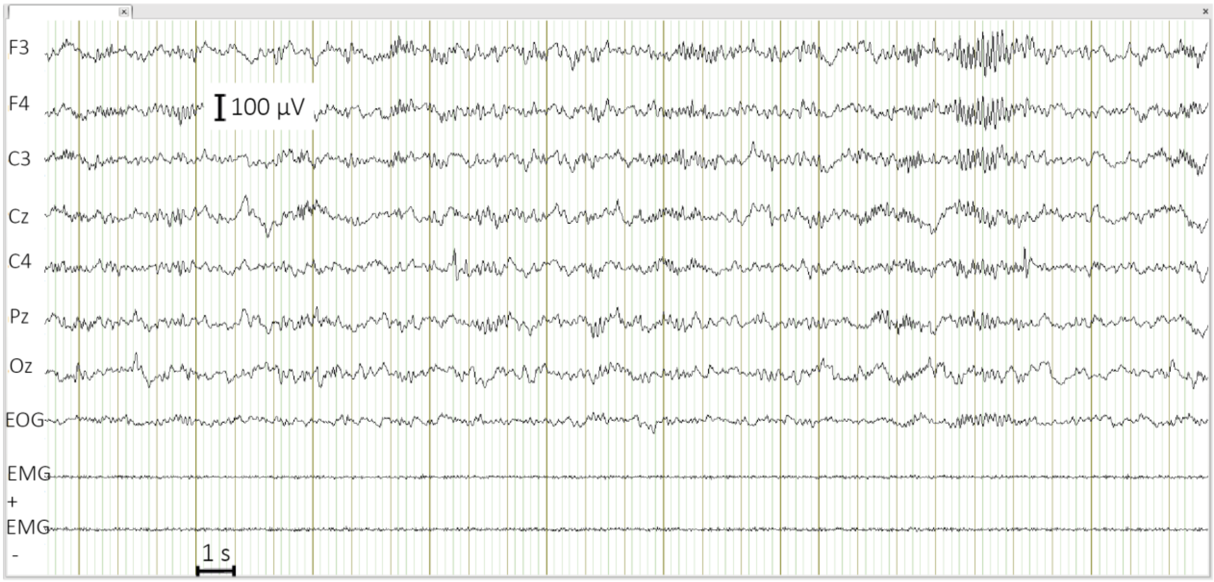


**D**

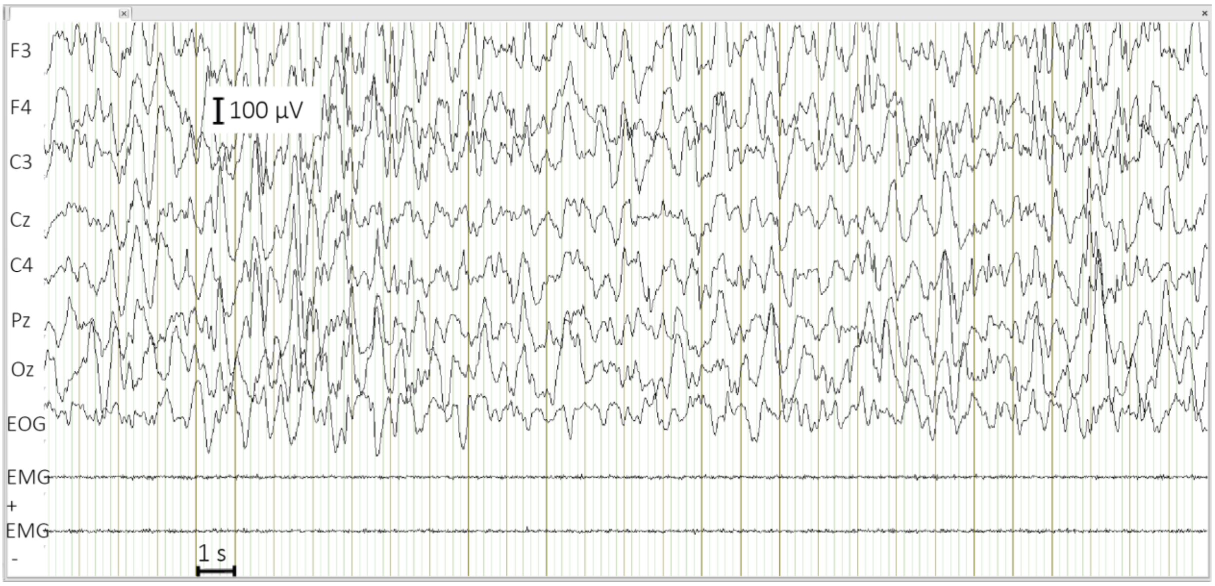


*Figure S2.* Raw EEG traces of a 5-year-old participant showing **(A)** sleep spindles and **(B)** slow waves and of a 6-year-old participant displaying **(C)** sleep spindles **(D)** slow waves.

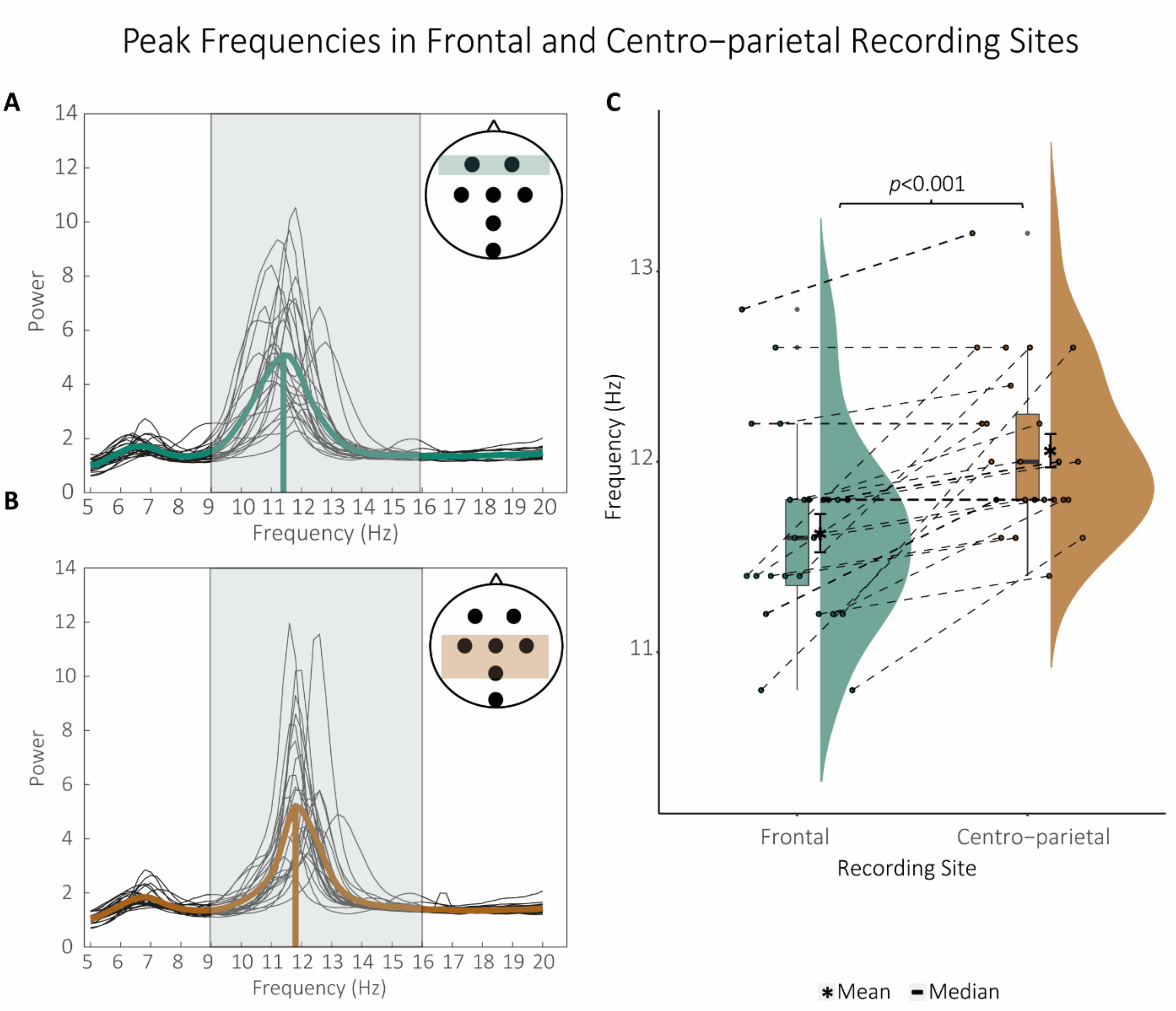


*Figure S3.* Averaged **(A)** frontal and **(B)** centro-parietal power spectra used to derive the **(C)** peak frequency for every participant. Bold coloured lines indicate the grand mean. **(C)** A Wilcoxon signed-rank test on the frontal and centro-parietal peak frequency revealed significantly slower peak frequencies frontal compared to centro-parietal electrodes ($Z$ = -3.82, $p$ < 0.001, $CI_{2.5, 97.5}$ [-4.20, -3.40]). Note that not every child showed two distinct peak frequencies in frontal and centro-parietal sites. Depicted here are measures derived during the learning night. Results for the baseline night show similar results.

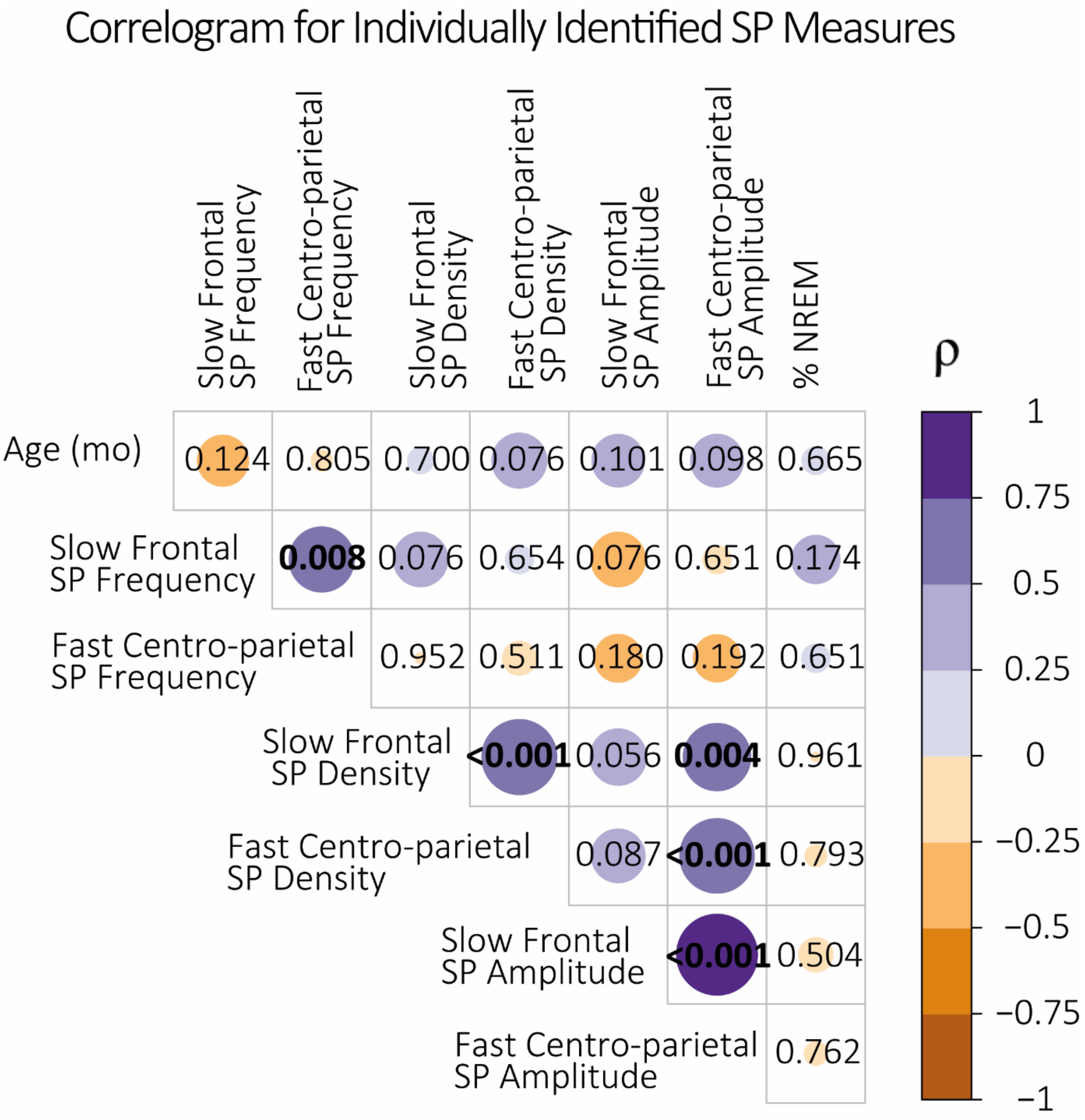


*Figure S4.* Correlations between measures of individually identified slow frontal and fast centro-parietal SPs during the learning night, age (in months), and the percentage of non-rapid eye movement sleep (% NREM) during the learning night. Values in the correlogram represent the *p*-values of Spearman's rank correlations. Colour coding indicates the magnitude of Spearman's rho. Significant correlations are highlighted in bold font.

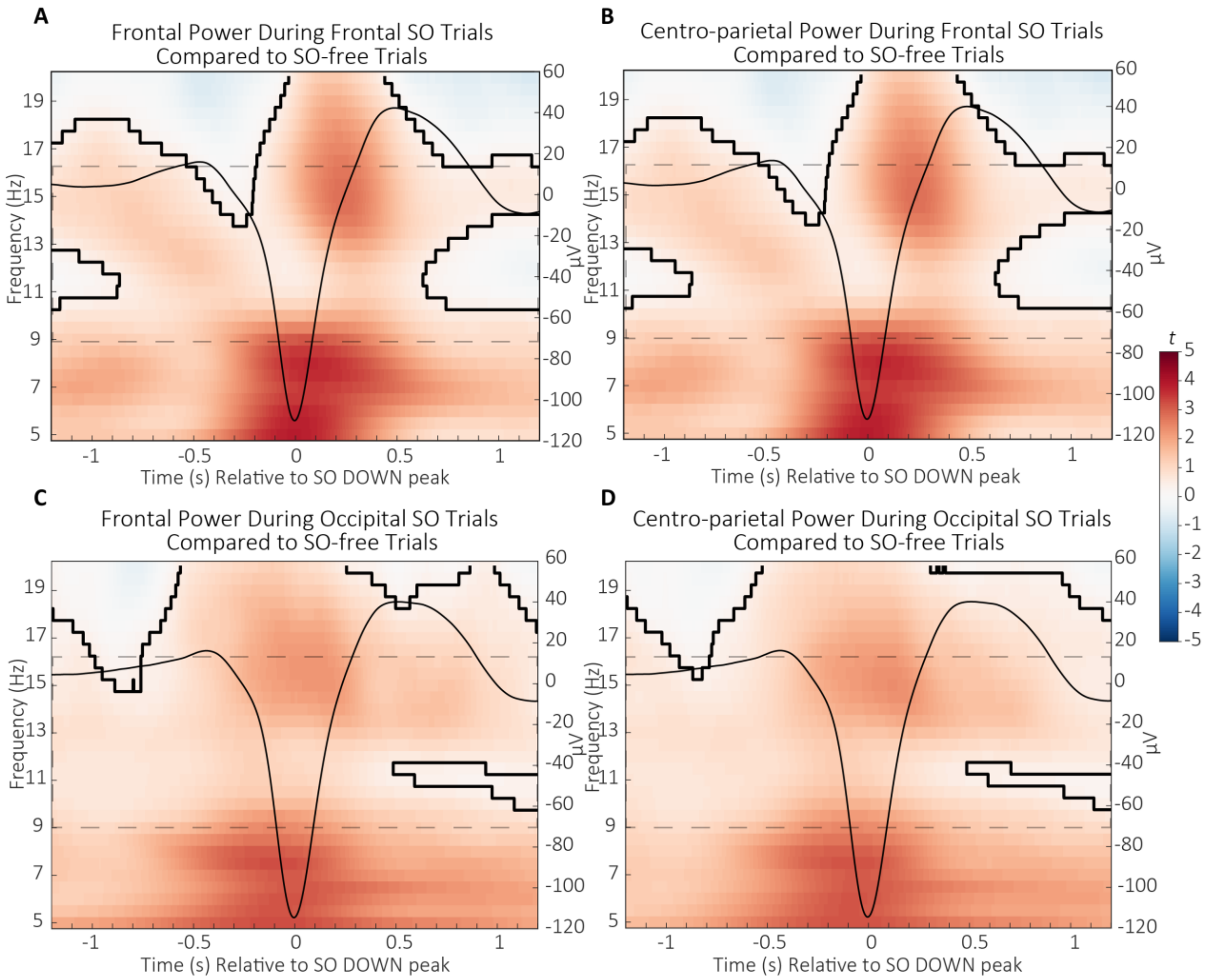


*Figure S5*. Differences in **(A)** frontal and **(B)** centro-parietal wavelet power during frontal SOs and **(C)** frontal and **(D)** centro-parietal wavelet power during occipital SOs compared to trials without SOs (*t*-score units) during the learning night. Significant clusters (cluster-based permutation test, cluster $\alpha < 0.05$, two-sided test) are outlined. The SP range is indicated by the reference window outlined in dashed lines. The average frontal/occipital SO is projected onto the power differences to illustrate their relation to the SO phase (scale in µV on the right side of each time-frequency plot).

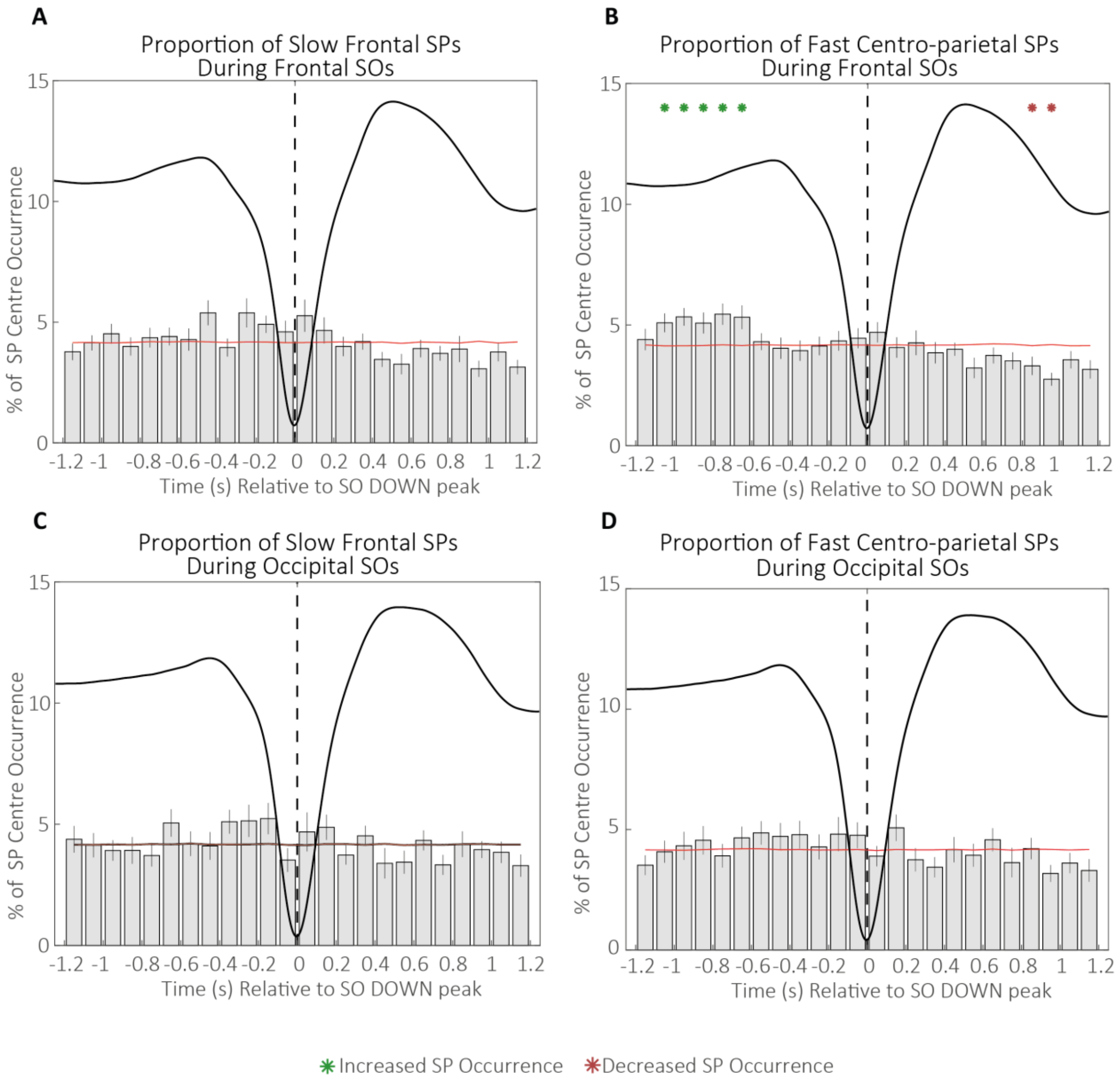


*Figure S6*. Percentage of individually identified **(A)** slow frontal and **(B)** fast cento-parietal SPs occurring within 100 ms bins during frontal SOs and of individually identified **(C)** slow frontal and **(D)** fast centro-parietal SPs occurring within 100 ms bins during occipital SOs during the learning night. Green asterisks indicate increased SP occurrence (positive cluster, cluster $\alpha$ < 0.05, two-sided test) and red asterisks indicate decreased SP occurrence (negative cluster, cluster $\alpha$ < 0.05, two-sided test) compared to random occurrence (black horizontal line with standard error of the mean indicated in red). Error bars represent standard errors. The dashed vertical line represents the SO DOWN peak. Averaged frontal and occipital SOs are depicted in black.

Power Spectra during Trials with “High” SPs

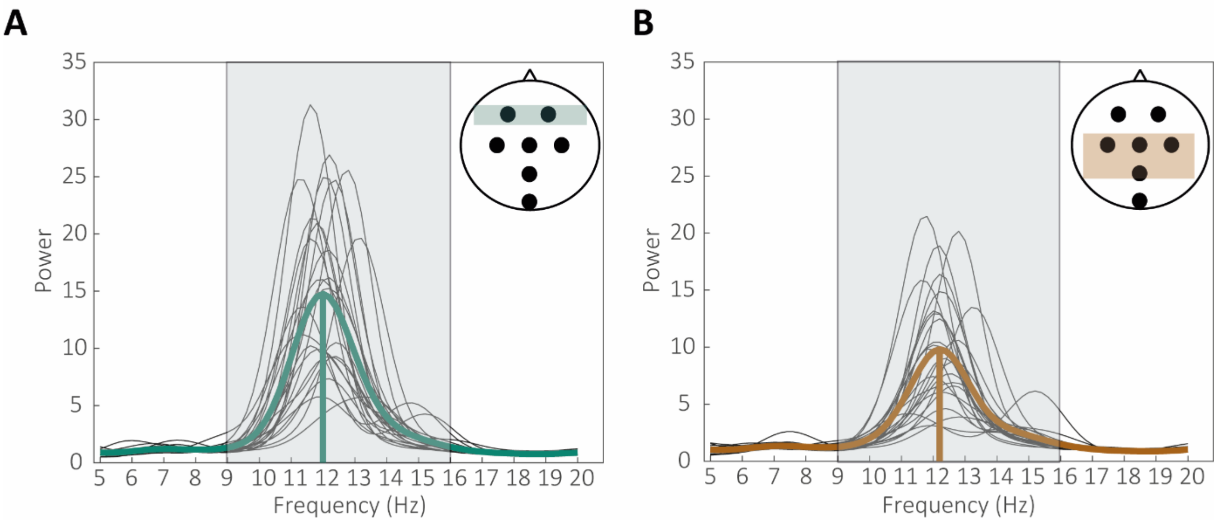


*Figure S7*. Averaged **(A)** frontal and **(B)** centro-parietal power spectra during trials in which “high” SPs were detected. Depicted here are measures derived during the learning night. Results for the baseline night are comparable.

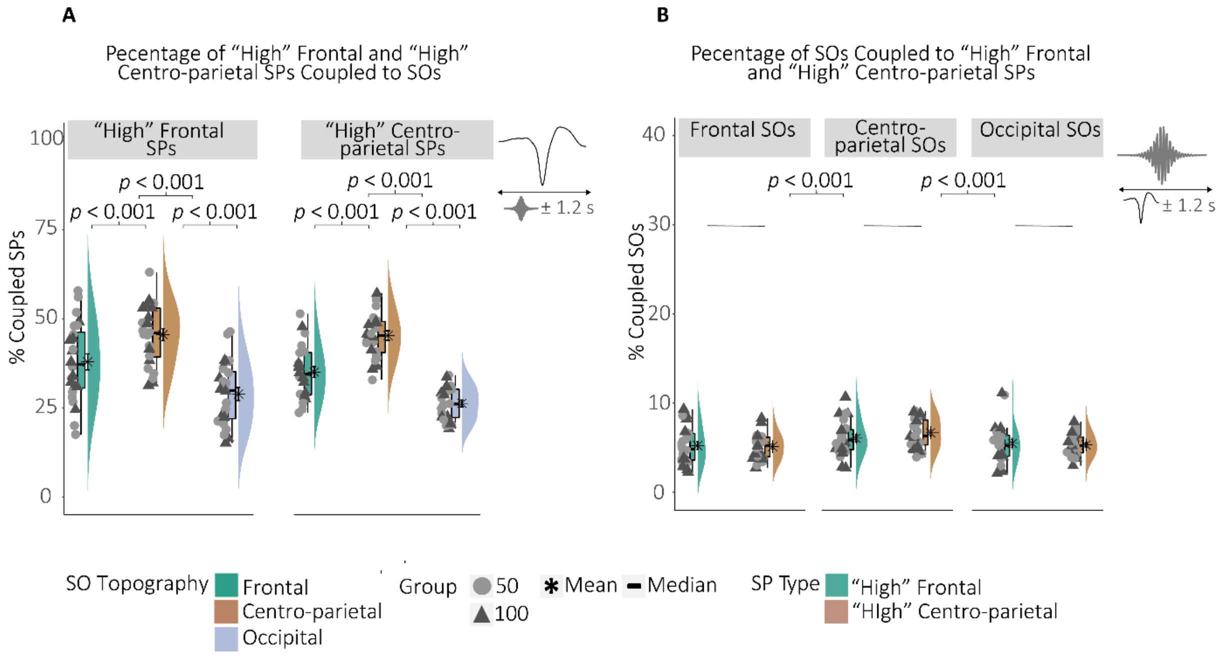


*Figure S8.* **(A)** Percentage of "high" frontal and "high" centro-parietal SPs co-occurring with frontal, centro-parietal, and occipital SOs during the learning night. *P*-values represent the results from the post-hoc Wilcoxon signed- rank tests from the ANOVA in Table S6. **(B)** Percentage of frontal, centro-parietal, and occipital SOs co-occurring with "high" frontal and "high" centro-parietal SPs. *P*-values represent the results from the post-hoc Wilcoxon signed-rank tests on the main effect "SO Topography", comparing centro-parietal SO DOWN peak co-occurrence with "high" SPs against frontal and occipital SO DOWN peak co-occurrence with "high" SPs.

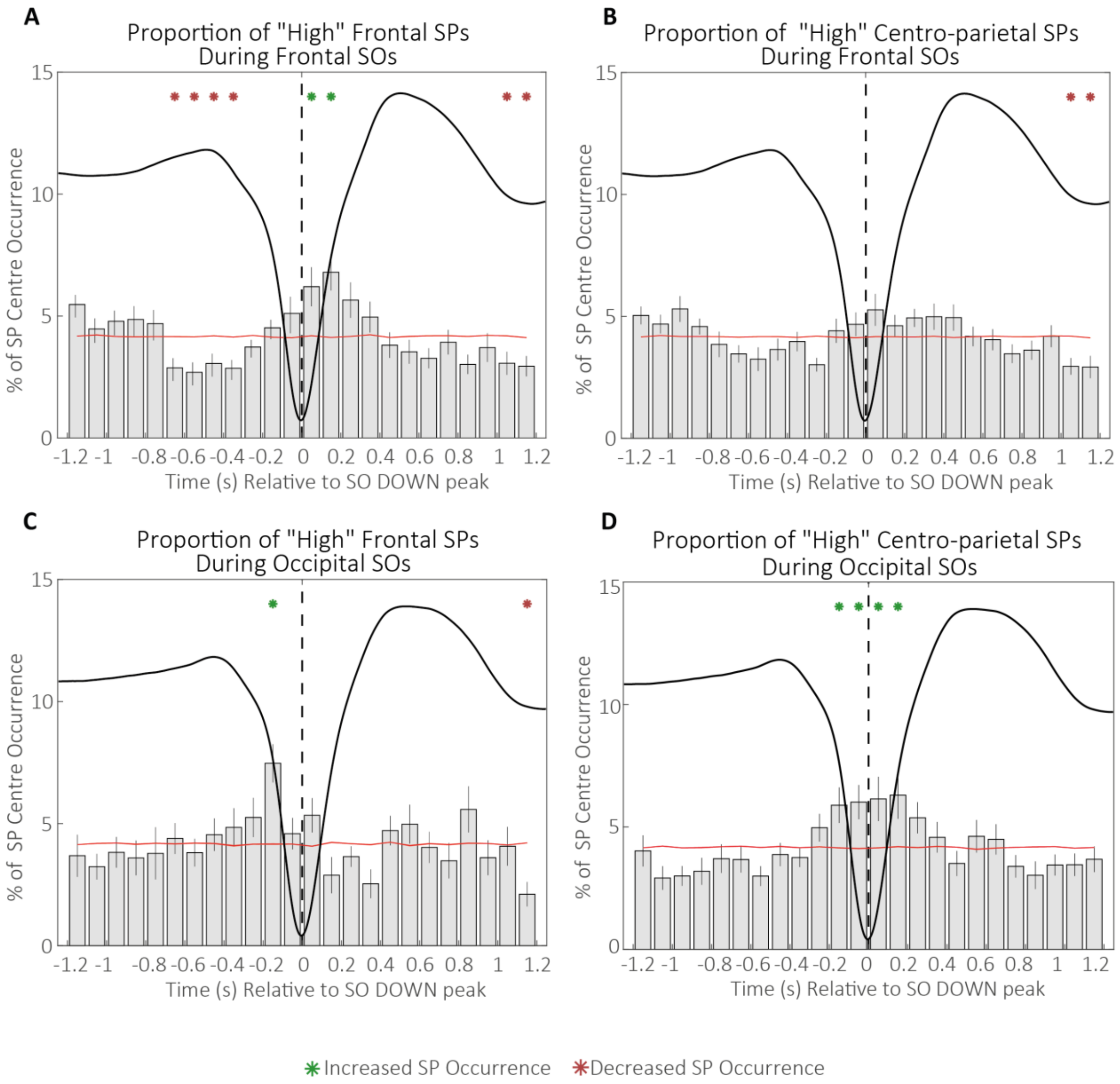


*Figure S9.* Percentage of "high" **(A)** frontal and **(B)** centro-parietal SPs occurring within 100 ms bins during frontal SOs and of "high" **(C)** frontal and **(D)** centro-parietal SPs occurring within 100 ms bins during occipital SOs during the learning night. Green asterisks indicate increased SP occurrence (positive cluster, cluster $\alpha < 0.05$, two-sided test) and red asterisks indicate decreased SP occurrence (negative cluster, cluster $\alpha < 0.05$, two-sided test) compared to random occurrence (black horizontal line with standard error of the mean indicated in red). Error bars represent standard errors. The dashed vertical line represents the SO DOWN peak. Averaged frontal and occipital SOs are depicted in black.

#### Correlograms for the Association between Indicators of SP-SO Coupling and Memory Consolidation

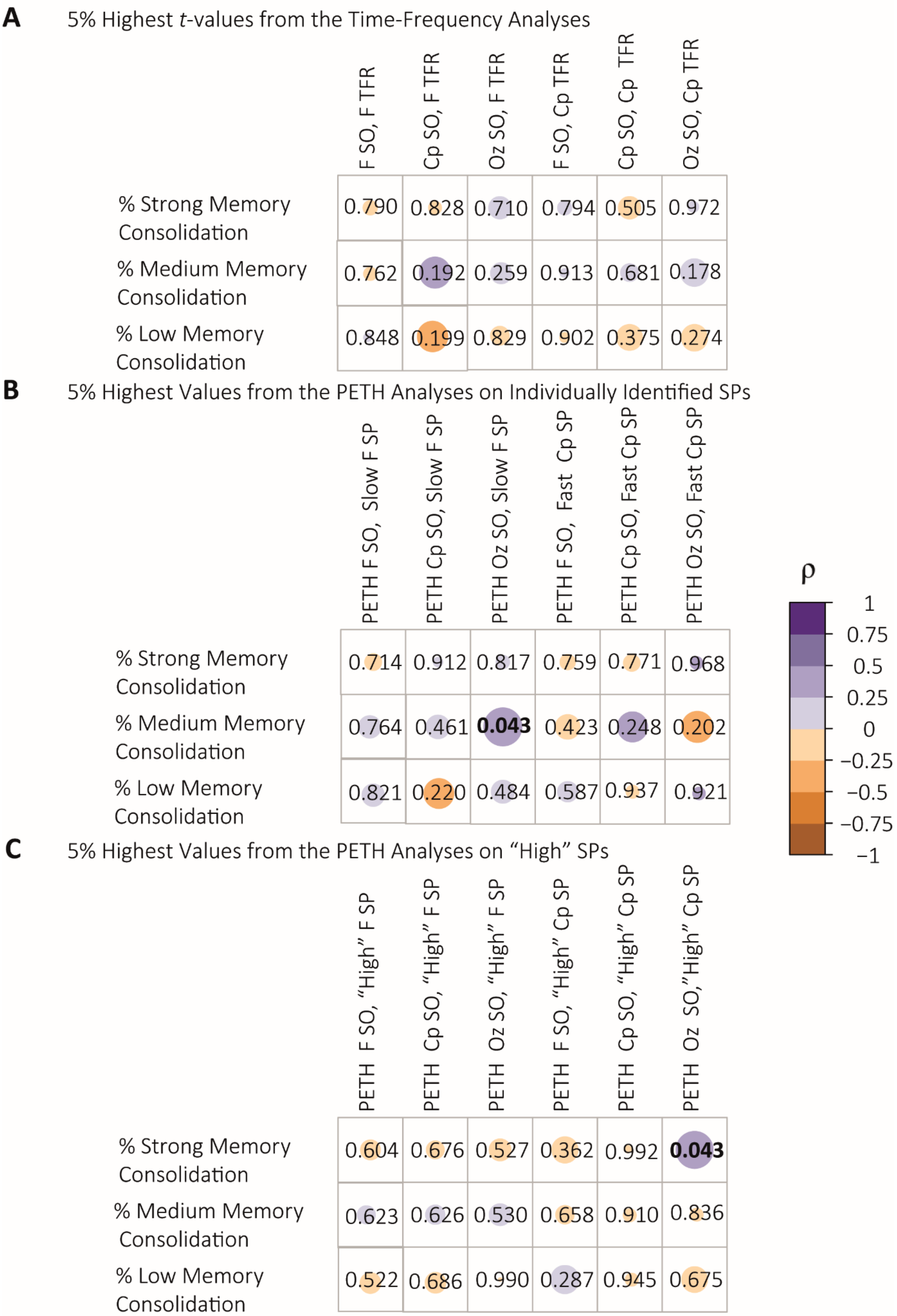

*Figure S10*. Correlations between indicators of SP-SO coupling during the learning night and measures of memory consolidation for items of varying quality **(A)** 5 % highest *t*-values from the group-level contrast of the time-frequency analyses. **(B)** 5% bins including the highest values of individually identified slow frontal and fast centro-parietal SP occurrence during SOs. **(C)** 5% bins including the highest values of “high” frontal and centro-parietal SP occurrence during SOs. Values in the correlograms represent the *p*-values of Spearman’s rank correlations. Colour coding indicates the magnitude of Spearman’s rho. Significant correlations are highlighted in bold font. Note: F indicates frontal, Cp indicates centro-parietal, and Oz indicates occipital topographical sites.
